# Prenatal fentanyl and Δ^9^-tetrahydrocannabinol exposure disrupt placental function and fetal growth in a mouse model of multidrug use

**DOI:** 10.64898/2026.09.17.752392

**Authors:** Yusmaris Cariaco, Nikita Larionov, Abolfazl Nik-Akhtar, Jessica Pudwell, Kira King, Laura Gaudet, Shannon Bainbridge

## Abstract

Opioid and cannabis co-use during pregnancy is increasingly common, yet the placental mechanisms linking combined exposure to adverse fetal outcomes remain poorly defined. Using a controlled mouse model of gestational drug exposure, we tested how fentanyl, Δ^9^-tetrahydrocannabinol (THC), or their combination altered placental structure, immune signaling, and gene expression and how these changes related to fetal growth. Drug exposure produced fetal growth restriction and reduced placental efficiency, with the greatest impairment in the combined fentanyl+THC group. Placental alterations were detectable by mid-gestation, when fentanyl exposure increased nucleated red blood cells within the labyrinth, consistent with hypoxic stress. By term, placentas showed compartment-specific remodeling, with THC selectively expanding the labyrinth and combined fentanyl+THC exposure increasing decidual area. Labyrinth composition and proliferative activity were altered, ultrastructural analysis revealed disruption of the maternal-fetal exchange interface, and placental interleukin-10 and interferon-β levels were reduced across exposure groups. Transcriptomic analyses identified suppression of innate immune and antiviral defense pathways together with treatment-specific stress responses, and integration of placental gene expression with fetal weight showed coordinated repression of vascular and developmental regulators and activation of hypoxia- and metabolic stress-associated genes. These findings identify the placenta as a key mediator of adverse fetal outcomes associated with prenatal polysubstance exposure.

**Significance Statement:** Opioids and cannabis are increasingly used together during pregnancy, but how this combination harms the developing fetus is unclear. Using a mouse model of chronic fentanyl and THC exposure, we show that combined exposure disrupts placental structure, suppresses immune signaling, and reprograms gene networks linked to fetal growth. These placental abnormalities appear before or alongside fetal growth restriction and are most severe with combined exposure. Our findings identify the placenta as a key biological target of prenatal polysubstance exposure and provide a mechanistic framework for understanding why opioid-cannabis co-use may worsen pregnancy outcomes.

## Introduction

Opioid use during pregnancy represents a critical and escalating dimension of the opioid epidemic, with profound implications for maternal and neonatal health. Across North America, reported prevalence estimates of opioid exposure during pregnancy range from approximately 1 % to 7 %, with substantial regional variability(1–6). Although some women reduce or discontinue opioid use during pregnancy, others continue or escalate use, often reflecting severe dependence(7). Additionally, opioid use disorder (OUD) at delivery has more than quadrupled between 1999 and 2014(8, 9), underscoring the urgency of addressing this crisis.

Although methadone and buprenorphine remain essential treatments for OUD, illicit fentanyl accounts for a disproportionate share of opioid-related morbidity and mortality(10). This shift is reflected in surveillance data showing that fentanyl-related overdose mortality among pregnant and postpartum individuals increased approximately 1.7-fold between 2017 and 2020(11). These epidemiological trends are mirrored in neonatal outcomes. In a recent clinical cohort of opioid-exposed pregnancies, nearly half of neonates had in-utero fentanyl exposure, which was associated with markedly greater severity of neonatal opioid withdrawal syndrome (NOWS) and significantly longer hospital stays(12). Together, these findings underscore the exceptional potency of fast-acting synthetic opioids such as fentanyl and their growing impact on perinatal health.

The complexity of opioid use disorder (OUD) during pregnancy is further compounded by the high prevalence of polysubstance use. Co-occurring use of alcohol, tobacco, and illicit drugs is common among pregnant individuals with opioid exposure and substantially amplifies risks for both maternal and fetal health(13–15). In clinical and population-based cohorts, non-opioid substance co-exposure is frequent, with cannabis emerging as the most common non-opioid drug, followed by benzodiazepines, amphetamines, cocaine, and other substances(16). The increasing normalization of cannabis use during pregnancy —particularly in regions with expanding legalization— introduces additional concern. Prenatal cannabis use varies by jurisdiction and measurement approach, ranging from ∼1–2 % in Ontario (2012–2017) to ∼7–8 % past-30-day use among U.S. individuals in early pregnancy, based on national survey analyses(9, 17, 18). Reported motivations for prenatal cannabis use frequently include self-management of pregnancy-related symptoms, such as nausea, anxiety, or stress(19–22). Within the context of opioid use disorder, this pattern of polysubstance use is particularly concerning given growing evidence that combined prenatal exposure to opioids and cannabis is associated with adverse birth and neurodevelopmental outcomes, including preterm birth, small size for gestational age, low birth weight, and neonatal intensive care unit (NICU) admission(23). Population-based analyses further demonstrate that maternal co-use of opioids and marijuana during pregnancy is associated with increased odds of prematurity and low birth weight compared with opioid exposure alone(24). Importantly, both fentanyl Δ^9^-tetrahydrocannabinol (THC) — the main psychoactive compound in cannabis— have been shown to cross the placenta and accumulate in the fetal brain, underscoring the heightened vulnerability of the developing nervous system to combined in-utero drug exposures(25–27).

Despite mounting epidemiologic and mechanistic evidence linking prenatal exposure to opioids or cannabis individually to adverse birth and neurodevelopmental outcomes(28–33), the placental mechanisms underlying combined exposure remain poorly understood. Prior studies demonstrate that opioids and cannabinoids each independently disrupt fetal growth trajectories(30, 34), neurodevelopment(35, 36), endocrine signaling(37), and metabolic programming(38); however, these investigations have largely examined single-drug exposure in isolation and provide limited insight into potential interactive or synergistic effects during pregnancy. The placenta plays a central role in regulating fetal growth, oxygen and nutrient exchange, immune tolerance, and endocrine signaling, yet the extent to which opioid and cannabinoid co-exposure perturbs placental structure, vascular organization, immune–cell dynamics, or functional capacity across gestation remains poorly defined.

To address these critical gaps, we employed a controlled mouse model to systematically evaluate the effects of fentanyl, THC, and their combined exposure on maternal metabolism, placental structure and function, and fetal growth and to determine how these exposures alter placental immune signaling, vascular organization, and fetal growth across gestation.

## Methods

### Drugs

THC was obtained from Protonofy (Product # PR1002-THC) and initially dissolved in 100% ethanol. This solution was then combined with Cremophor EL (MedChemExpress, HY-Y-1890) and saline in a ratio of 1:1:18 (THC/ethanol: Cremophor EL: saline). The matched vehicle contained ethanol, Cremophor EL, and saline in the same proportions but without THC. Animals not receiving THC were administered an equivalent volume of this vehicle according to the same schedule.

Exemptions for the use of controlled substances for scientific purposes were granted by Health Canada (protocol numbers 54319.07.22AMD & 58028.08.24).

### Mice

All procedures were approved by the ACVS Ethics Review Board (protocol #3895) and performed in the animal facilities of the University of Ottawa. Pregnant CD1 mice (Charles River Laboratories) were randomly assigned to one of four experimental groups: (1) Control (vehicle; Ctrl); (2) fentanyl (0.3 mg/kg/day; Fen); (3) THC (3 mg/kg/day); and (4) combined fentanyl + THC exposure (FenTHC). Treatments were administered once daily by subcutaneous (s.c.) injection as described. Mice, maintained under a 12-hour light/dark cycle with ad libitum access to food and water at the University of Ottawa Animal Care Veterinary Services (ACVS) facilities, were administered fentanyl for 2 weeks prior to mating to model chronic opioid use. Δ-9-THC treatment began on embryonic day (E)6.5 to mitigate the risk of embryo loss associated with earlier exposure. The day of vaginal plug detection was designated as embryonic day E0.5. At euthanasia on E18.5, litter size, embryo/fetus viability, and fetal and placental weights were recorded. Placentas were bisected, with one half fixed in formalin for histological processing and the other half flash-frozen for protein/RNA analysis. Sex was determined by *Sry* genotyping of fetal tail tissue, and placentas were pooled by fetal sex for subsequent analyses.

Fetal viability was assessed by classifying each conceptus at the time of collection into one of four categories. Early resorptions were identified as small, dark, avascular remnants, typically lacking visible placental development, suggesting that embryonic loss occurred prior to placentation. Late resorptions were defined by the presence of a formed placenta with an associated degenerated or avascular conceptus, often appearing as a markedly smaller, pale, or hemorrhagic mass, indicating fetal demise after placental formation. Non-viable fetuses were fully formed with corresponding placental development but exhibited clear signs of death, including pallor, hemorrhagic discoloration, and absence of responsiveness to tactile stimulation. Viable fetuses were characterized by anatomically complete fetal and placental structures, a healthy external appearance, and a positive response to a toe-pinch stimulus, even in cases of apparent growth restriction.

### Fetal sex determination

To determine fetal sex, DNA was extracted from fetal tail samples using Extract-N-Amp™ Tissue Extraction Solution (Sigma-Aldrich, E7526), followed by neutralization with DNA Neutralization Solution (Sigma-Aldrich, N3910). Genomic DNA was subjected to Polymerase Chain Reaction (PCR) using Phire™ Tissue Direct PCR Master Mix.

SRY gene amplification was achieved with the following primers: forward primer 5′-TTGTCTAGAGAGCATGGAGGGCCATGTCAA-3′ and reverse primer 5′-CCACTCCTCTGTGACACTTTAGCCCTCCGA-3′.

### Indirect calorimetry and metabolic phenotyping

Metabolic parameters were assessed using Oxymax chambers of the Comprehensive Lab Animal Monitoring System (CLAMS; Columbus Instruments, Columbus, OH, USA). Mice were housed individually in the metabolic chambers for a total of 96 hours with continuous data collection throughout the entire period. The first 72 hours were used for habituation, and data from the final 24 hours were used for analysis.

Animals were maintained on a standard 12-hour light-dark cycle with ad libitum access to food and water throughout the testing period. The chow provided during CLAMS assessment was the same formulation used in their regular housing, supplied in powdered form to accommodate the automated feeding system. Food intake was monitored via automated feeders integrated within the CLAMS system, allowing precise quantification of intake patterns. Energy expenditure (EE) was calculated using indirect calorimetry based on the recorded VO₂ and VCO₂ values according to the standard Weir equation. Locomotor activity was assessed as ambulatory movement, quantified by horizontal (X-axis) infrared beam breaks within the CLAMS chambers.

Body composition was measured prior to CLAMS assessment using EchoMRI (EchoMRI LLC, Houston, TX, USA) to determine lean mass and fat mass, and metabolic parameters were normalized to lean mass. All measurements were conducted within an environmental chamber maintained at a constant temperature of 28°C and relative humidity of approximately 40–60%. Data were analyzed using Oxymax software (Columbus Instruments).

### Electron microscopy

E18.5 placentas were fixed in 0.1 M sodium cacodylate buffer (pH 7.4) for 24 h. Fixed tissues were processed by the uOttawa Transmission Electron Microscopy (TEM) Core Facility and sectioned into 50 µm free-floating slices using a vibratome. Regions of interest were excised from these sections and mounted onto resin blocks using cyanoacrylate adhesive. Ultrathin sections (140 nm) were then cut in ribbons using an ultramicrotome and collected on 100-mesh copper grids. The placental labyrinth layer was imaged at 20,000× magnification using a JEOL JEM-1400Flash transmission electron microscope operated at 120 kV and equipped with a Gatan OneView 4K CMOS digital camera and a LaB₆ filament to enhance image contrast.

### Histological processing

Formalin-fixed placentas were processed by the uOttawa Louise Pelletier Histology Core Facility for paraffin embedding, sectioning, hematoxylin and eosin (H&E) staining, and immunohistochemistry. Immunohistochemical staining was performed using the Leica Bond™ automated staining platform following a modified version of Protocol F, in which the post-primary step was omitted for rabbit antibodies on mouse tissue.

Heat-mediated antigen retrieval was carried out in sodium citrate buffer (pH 6.0; Epitope Retrieval Solution 1) for 20 min. Sections were then incubated with primary antibodies for 30 min at room temperature: rabbit anti-CD31 (1:100, Novus Biologicals, NB100-2284), rabbit anti-Ki67 (SP6 clone, 1:200, Abcam, ab16667), and rabbit anti-CD68 (1:1500, Abcam, ab125212). Detection was achieved using an HRP-conjugated compact polymer detection system, followed by visualization with DAB chromogen.

Slides were counterstained with hematoxylin, mounted, and cover slipped. Whole-slide images were acquired using a Zeiss Axio Scan Z.1 slide scanner.

Image analysis was performed using QuPath (v0.5.0) (39). Vascular compartments within the labyrinth were segmented using supervised pixel classification trained to distinguish CD31⁺ endothelial structures from surrounding placental tissue and maternal blood spaces at the late gestation timepoint. For mid-gestation samples, classifiers were trained to identify DBA⁺ fetal capillaries and to separate these from cytokeratin-positive trophoblast and cytokeratin-lined maternal blood spaces based on multiplex signal patterns (DBA, cytokeratin, and DAPI). These classifications enabled automated quantification of vascular area within annotated labyrinth regions.

Macrophage (CD68⁺) and proliferative (Ki67⁺) cells were quantified across decidual and whole placental sections, respectively, using automated detection workflows, with positive events normalized to the corresponding annotated tissue area.

Placental layer composition was assessed through manual annotation of histological compartments—including the decidua, junctional zone, and labyrinth—based on established morphologic criteria, from which absolute regional areas (µm²) were calculated.

Nucleated red blood cells (nRBCs) were quantified within the labyrinth following whole-cell detection across annotated regions. An object classifier was trained to identify erythroid cells based on nuclear morphology, hematoxylin intensity, and cell size features, and classified nRBCs were expressed relative to the total number of detected cells within the labyrinth.

### ELISA

Placental protein extracts were prepared by homogenizing placental tissue in phosphate-buffered saline (PBS) containing protease inhibitors (cOmplete™, Roche #4693159001). For each dam, placentas from fetuses of the same sex were pooled prior to homogenization using a bead mill homogenizer (Fisherbrand, 15-340-163). The homogenates were centrifuged at 16,000 × g for 10 minutes. Protein concentrations for normalization were determined using the DC Protein Assay Kit (Bio-Rad, 5000112).

Subsequent measurements of IFN-β (Biolegend, 439407), IL-10 (Biolegend, 431417), 5-HT (Abcam, ab133053), and folic acid (Aviva Systems Biology, OKEH02550) were performed according to the manufacturer’s instructions.

### RNA sequencing

Ten samples per exposure group and control were included, comprising five male and five female pooled placental samples per group. Within each dam, placentas from fetuses of the same sex were pooled prior to RNA extraction. Total RNA was extracted using TRIzol reagent (Life Technologies, #15596026) according to the manufacturer’s protocol. RNA quality was assessed at the Centre for Applied Genomics (The Hospital for Sick Children, Toronto) using an Agilent Bioanalyzer, confirming RNA integrity numbers (RIN) > 7. Stranded poly(A) mRNA libraries were prepared using the NEBNext® Ultra™ II Directional mRNA Library Prep Kit, and sequencing was performed on a NovaSeq 6000 platform with an S4 flow cell, generating 33–42 million 150 bp paired-end reads per sample. Untrimmed FASTQ files were processed on the Galaxy platform. Reads were preprocessed using fastp for quality control, adapter removal, quality filtering, and trimming. Processed reads were aligned to the mouse reference genome using HISAT2, and gene-level counts were generated using featureCounts.

### Statistical Analysis

Statistical analyses were performed using GraphPad Prism 10.0 and RStudio 2025.05.0. Data are presented as mean ± SEM, with significance defined as p < 0.05. The dam was considered the experimental unit for litter-derived measurements.

Individual fetuses or placentas were treated as observational units, with fetus–placenta measurements within each litter treated as repeated measures. Values were averaged within each litter prior to statistical analysis. Main effects were analyzed using a two-way factorial model to evaluate the independent and combined effects of fentanyl and THC. Post hoc comparisons were performed using uncorrected Fisher’s least significant difference (LSD) test. For sex-stratified analyses, drug exposure and fetal sex were included as factors, using Dunnett’s post hoc test was to compare treatment groups with controls. Time-course metabolic data were analyzed by two-way ANOVA with treatment and time as factors.

RNA-seq differential expression was performed using an edgeR-based pipeline with the design formula ∼ 0 + DrugExposure + Sex, and contrasts comparing each exposure to controls. Sex-specific responses were tested using an interaction model (DrugExposure × Sex). Genes were retained if ≥10 total counts and ≥2 CPM in ≥5 samples, and significance was defined as Benjamini–Hochberg FDR < 0.05.

Associations between placental gene expression and fetal weight were assessed using sex-adjusted partial Pearson correlations (ppcor) with FDR correction. Differential expression was visualized using ggplot2 volcano plots (FDR < 0.05, |log₂FC| > 1).

Functional enrichment was performed using WebGestalt (GO over-representation analysis) and clusterProfiler (GSEA). Additional visualizations were generated using pheatmap, UpSetR, corrplot, and ggplot2.

## Results

### Pre-pregnancy fentanyl and THC exposure alters baseline metabolic parameters in non-pregnant females

Using a mouse model of pre-gestational drug exposure to mimic chronic opioid use before pregnancy (Fig. 1A), we first confirmed that fentanyl exposure in females designated for timed pregnancies did not alter body weight during the two weeks preceding mating (Fig. 1B). However, unchanged body weight does not rule out underlying alterations in metabolic function. Because pregnancy itself profoundly reshapes maternal metabolism, distinguishing the direct metabolic effects of fentanyl and THC from those that arise secondary to gestation is essential. Since our gestational exposure paradigm does not include pre-pregnancy THC or combined fentanyl–THC exposure, we established a baseline in a separate cohort of non-pregnant females exposed to fentanyl (Fen), THC, or the combination (FenTHC) for nine days, and assessed metabolic function using the CLAMS metabolic system (Fig. 1A). This baseline framework allowed us to interpret later gestational metabolic outcomes in the context of drug-specific effects.

**Figure 1.**
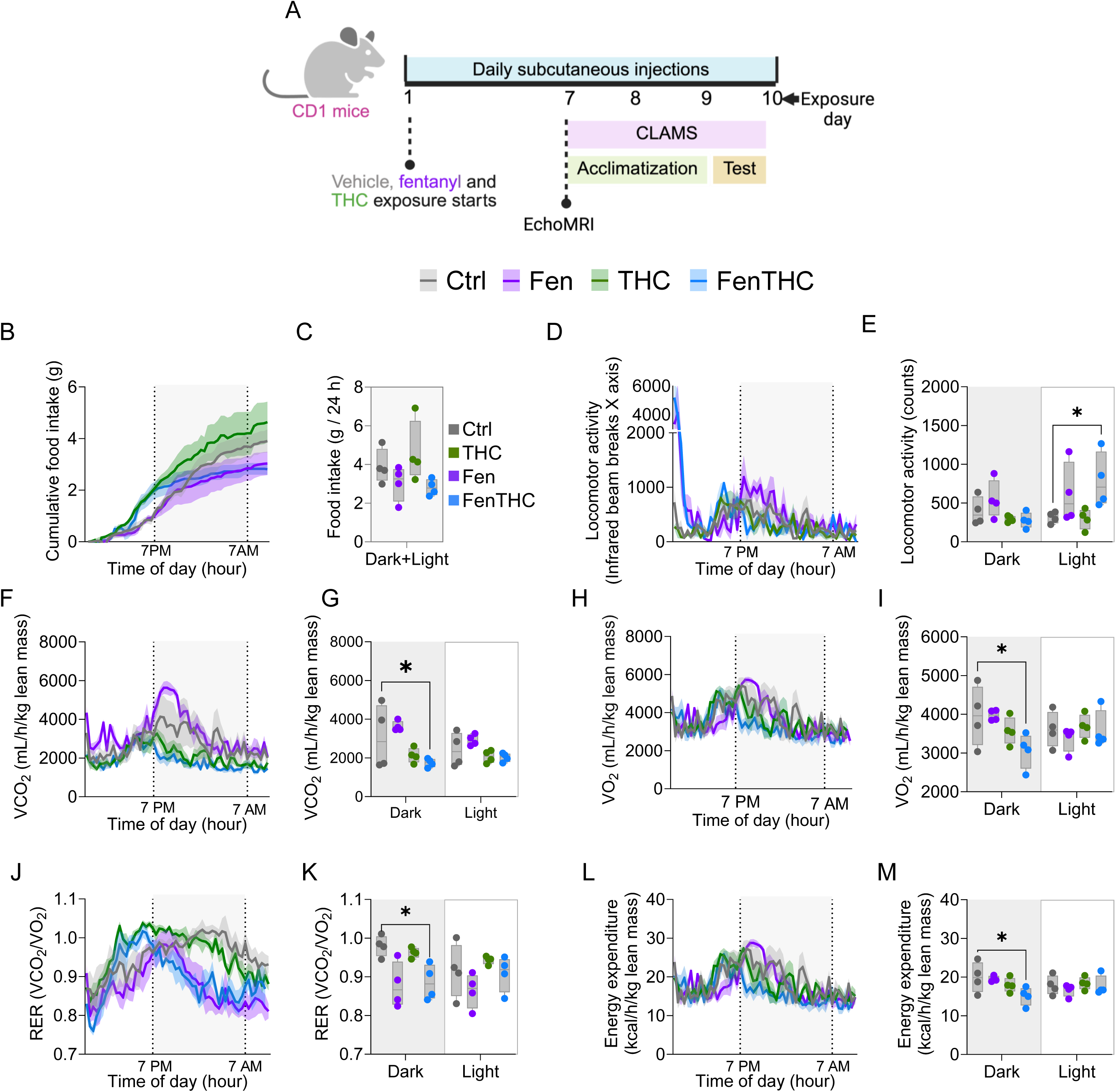
Pre-pregnancy fentanyl and Δ^9^-THC exposure alters baseline metabolic parameters in non-pregnant females. (A) Experimental timeline for pre-gestational drug exposure and indirect calorimetry in non-pregnant CD1 females. (B) Percent change in body weight across the two-week of chronic opioid exposure period (control n=20 vs fentanyl, n=20). (C) Cumulative food intake across the final 24 h of Comprehensive Lab Animal Monitoring System (CLAMS) monitoring. (D) Total food intake over 24 h. (E) Ambulatory locomotor activity (infrared beam breaks) across the 24 h monitoring period. (F) Mean locomotor activity quantified by light and dark phases. (G–H) Carbon dioxide production (VCO_2_) over time (G) and averaged by phase (H). (I– J) Oxygen consumption (VO_2_) over time (I) and averaged by phase (J). (K–L) Respiratory exchange ratio (RER) over time (K) and averaged by phase (L). (M–N) Energy expenditure (EE) over time (M) and averaged by phase (N). For time-course panels, shaded background denotes the dark phase and dashed vertical lines indicate lights-off/on. VO_2_, VCO_2_, and EE are normalized to lean mass. CLAMS experiments had n=4/group, points represent individual mice and lines/bars show mean ± SEM. Statistical analysis was performed using two-way ANOVA followed by Dunnett’s post hoc test comparing treatment groups to controls. Ctrl, control; THC, Δ^9^-tetrahydrocannabinol; Fen, fentanyl; FenTHC, fentanyl + Δ^9^-tetrahydrocannabinol. *p < 0.05.

THC exposure produced a transient increase in food intake immediately preceding the dark phase, consistent with the onset of the mice’s peak activity period, with no differences in 24-hour cumulative food intake between treatment groups and controls, either during the two-week monitoring period or in the fentanyl-treated group (Fig. 1C– D, Supplementary Fig. 1). In contrast, fentanyl exposure was associated with a transient increase in locomotor activity during the light phase immediately following daily injections, an effect that reached significance when fentanyl was combined with THC (Fig. 1E–F). This acute hyperlocomotion is consistent with a short-term pharmacodynamic response to fentanyl that appears amplified by concurrent THC exposure.

Co-exposure to fentanyl and THC disrupted normal metabolic physiology during the dark phase. FenTHC-treated mice displayed significant reductions in VO₂ and VCO₂ (Fig. 1G–J), accompanied by a lower respiratory exchange ratio (RER) (Fig. 1K–L), indicating respiratory depression and a shift toward preferential fat oxidation, and a corresponding decrease in energy expenditure (Fig. 1M–N). Together, these findings show that combined opioid and cannabis exposure alters substrate utilization and suppresses metabolic rate in the absence of pregnancy, establishing an important metabolic baseline for interpreting subsequent gestational effects.

### Pregnancy buffers against metabolic disturbances induced by fentanyl and THC

We next evaluated how opioid and cannabis exposure influences maternal metabolism during pregnancy. Although fentanyl-exposed dams gained weight similarly to controls, mice exposed to THC alone or in combination with fentanyl exhibited reduced gestational weight gain beginning at E10.5, coinciding with early chorioallantoic placental development and indicating a THC-associated deviation from the expected trajectory of maternal weight gain (Fig. 2A).

**Figure 2.**
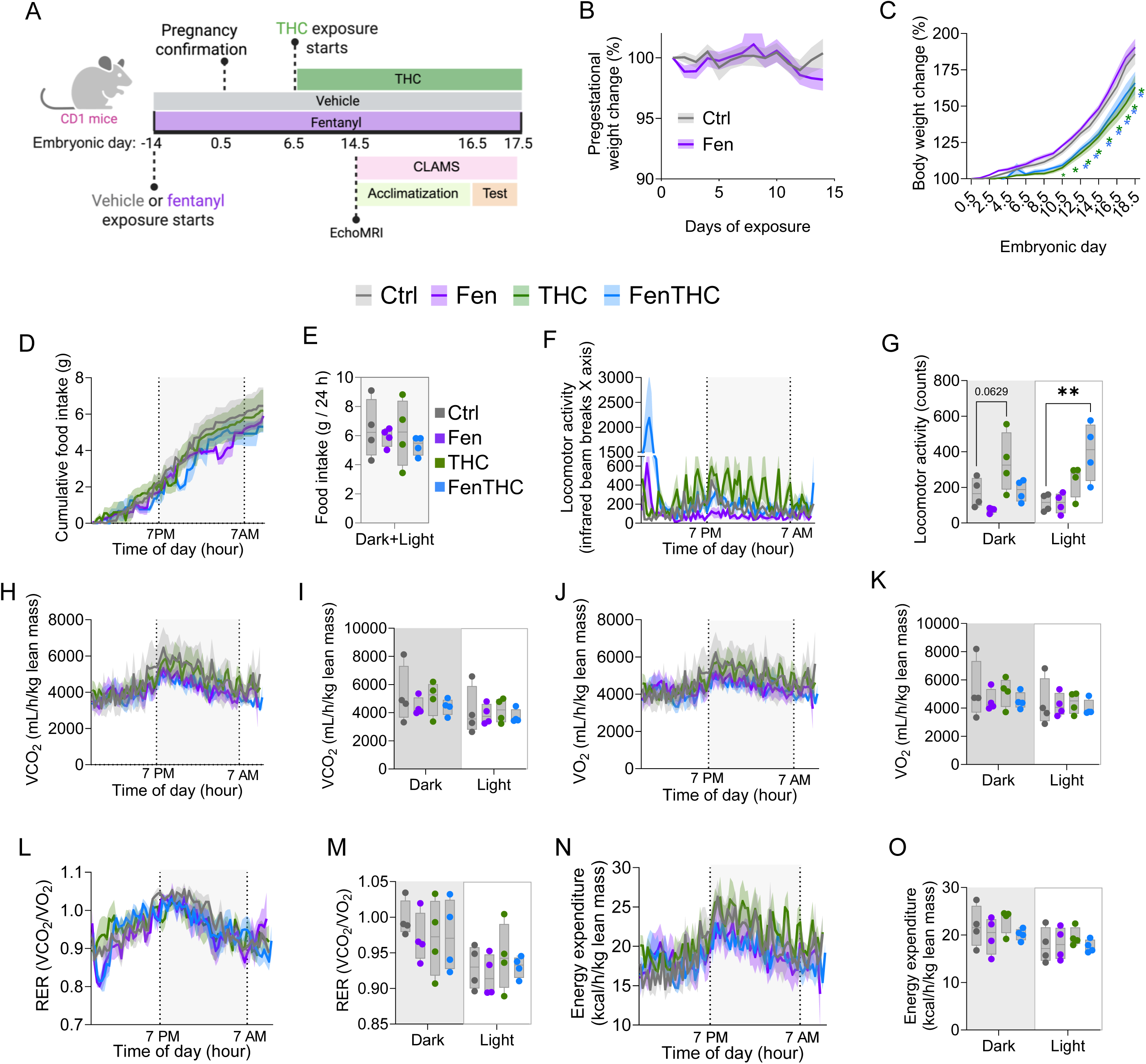
Pregnancy buffers against metabolic disturbances induced by fentanyl and Δ^9^-THC exposure. (A) Gestational body weight change (weight at E0.5 = 100%) across pregnancy in Ctrl-, THC-, fentanyl-, and FenTHC–exposed dams. Green and blue asterisks represent statistical differences at the given timepoints between THC and FenTHC vs Ctrl, respectively. (B) Experimental timeline for fentanyl exposure and gestational THC exposure (initiated at E6.5), with indirect calorimetry performed at E16.5. (C) Cumulative food intake across the final 24 h of Comprehensive Lab Animal Monitoring System (CLAMS) monitoring at E16.5. (D) Total food intake over 24 h. (E) Ambulatory locomotor activity (infrared beam breaks) across the 24 h monitoring period. (F) Mean locomotor activity quantified by light and dark phases. (G–H) Carbon dioxide production (VCO_2_) over time (G) and averaged by phase (H). (I–J) Oxygen consumption (VO_2_) over time (I) and averaged by phase (J). (K–L) Respiratory exchange ratio (RER) over time (K) and averaged by phase (L). (M–N) Energy expenditure (EE) over time (M) and averaged by phase (N). For time-course panels, shaded background denotes the dark phase and dashed vertical lines indicate lights-off/on. VO_2_, VCO_2_, and EE are normalized to lean mass. CLAMS experiments had n=4/group, points represent individual dams and lines/bars show mean ± SEM. Statistical analysis was performed using two-way ANOVA followed by Dunnett’s post hoc test comparing treatment groups to controls. Ctrl, control; THC, Δ^9^-tetrahydrocannabinol; Fen, fentanyl; FenTHC, fentanyl + Δ^9^-tetrahydrocannabinol. *: p < 0.05, **: p < 0.01, ***: p < 0.001, ****: p < 0.0001.

To determine whether these changes reflected altered maternal metabolic function, a separate cohort of pregnant females was exposed to fentanyl, THC, or their combination and assessed at E16.5 using the CLAMS metabolic system (Fig. 2B). Over the 24-hour monitoring period, cumulative food intake did not differ among groups (Fig. 2C–D). As observed in non-pregnant mice, fentanyl exposure induced a sharp, transient increase in locomotor activity immediately following injection during the light phase. This spike reflects an acute pharmacodynamic response to fentanyl that rapidly dissipates, followed by reduced activity for the remainder of the monitoring period. In contrast,

THC-treated mice did not display a pronounced post-injection peak, but instead showed a sustained elevation in locomotor activity throughout the dark phase. Because this increase is maintained over time, THC exposure results in a higher overall average locomotor activity during the dark phase. The combined FenTHC group exhibits features of both responses: an exaggerated immediate post-injection peak driven by fentanyl and a more persistent elevation in activity resembling the THC pattern. This distinction suggests that fentanyl primarily drives acute, short-lived hyperlocomotion, whereas THC modulates baseline arousal or activity state across the active period, leading to a greater cumulative locomotor output.

Measurements of VCO_2_, VO_2_, and energy expenditure showed largely overlapping profiles across treatment groups, with no evidence of respiratory suppression. Although the overall respiratory exchange ratio (RER) did not differ between groups, both Fen and FenTHC mice exhibited a significant transient reduction in RER immediately after injection and at the onset of the dark phase. This decrease indicates brief shifts toward greater reliance on fat oxidation, consistent with the more pronounced metabolic alterations observed in non-pregnant mice (Fig. 2G–N).

Collectively, these findings indicate that, despite clear effects on gestational weight gain, exposure to fentanyl, THC, or their combination during pregnancy induces only subtle changes in maternal metabolic physiology at E16.5. The primary detectable effects consist of transient behavioral responses and mild, time-restricted alterations in respiratory metabolism, suggesting that pregnancy buffers or stabilizes systemic metabolic homeostasis even in the presence of drug exposure.

### Fentanyl and THC exposure perturb placental cellular dynamics at mid-gestation

Given that gestational weight gain began to diverge by E10.5 in the THC and FenTHC groups, we asked whether fetal growth restriction is already detectable at this early window and whether it coincides with measurable changes in placental composition and organization. To address this, we profiled fetal and placental growth metrics and quantified key histologic and cellular features of the maternal–fetal interface at mid-gestation (E13.5) (Fig. 3A).

**Figure 3.**
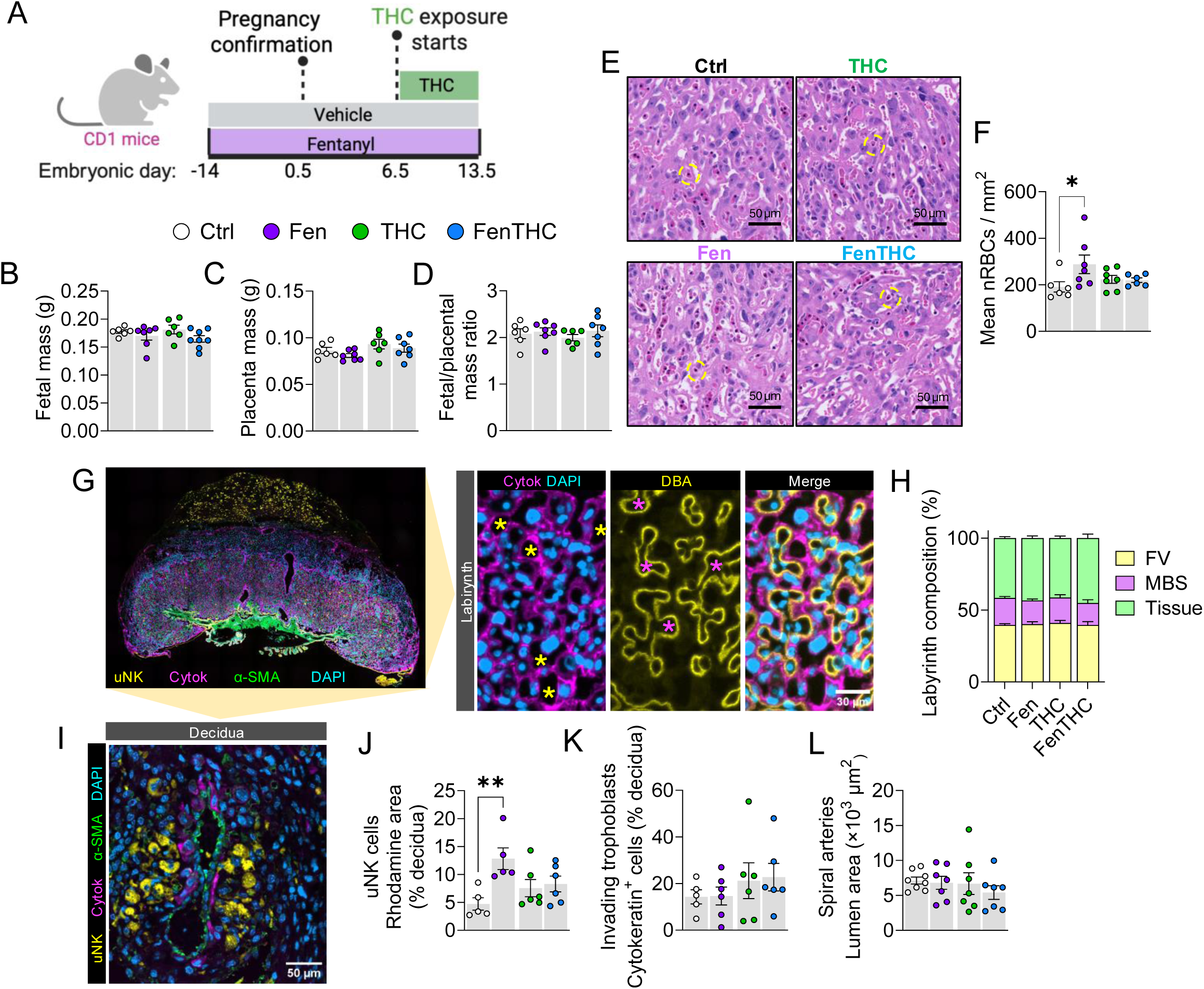
Fentanyl and Δ^9^-THC exposure induces fetal growth restriction and early placental alterations at mid-gestation. (A) Experimental design: mice were exposed or not to fentanyl two weeks before pregnancy confirmation at E0.5, fentanyl was administered continuously from E0.5 to E13.5, and THC exposure began at E6.5 and continued through E13.5, with vehicle-treated mice serving as controls. (B) Fetal weight at E13.5. (C) Placental weight at E13.5. (D) Placental efficiency (fetal/placental weight ratio) at E13.5. (E) Representative hematoxylin and eosin (H&E) images of the placental labyrinth layer from each exposure group. (F) Quantification of nucleated red blood cells (nRBCs) in the labyrinth (nRBCs/mm²), as an index of fetal/placental hypoxic stress. (G) Representative immunofluorescence images of whole placenta and labyrinth region showing cytokeratin (trophoblast; “Cytok”), DBA lectin signal (fetal blood vessels), and DAPI nuclear staining. (H) Labyrinth compartment composition quantified as the relative proportion of fetal vessels (FV), maternal blood spaces (MBS), and trophoblast tissue. (I) Representative decidual immunostaining highlighting uterine NK cells (uNK; DBA^+^ - yellow), invading trophoblasts (cytokeratin^+^ - pink), vascular smooth muscle (α-SMA - green), and nuclei (DAPI - blue). (J) Quantification of uNK abundance in the decidua (DBA/rhodamine area as % of decidual area). (K) Quantification of trophoblast invasion (cytokeratin^+^ cells as % of decidual area). (L) Spiral artery lumen area. Each point is the litter mean (one point per dam, n=5-7/group) and bars show mean ± SEM. Statistical analysis was performed using two-way ANOVA followed by uncorrected Fisher’s LSD post hoc test. Ctrl, control; THC, Δ^9^-tetrahydrocannabinol; Fen, fentanyl; FenTHC, fentanyl + Δ^9^-tetrahydrocannabinol. *: p < 0.05, **: p<0.01.

At E13.5, fetal size remained unchanged between FenTHC and control groups (Fig. 3B). Placental weight (Fig. 3C) and the fetal-to-placental weight ratio (Fig. 3D) were also unchanged, indicating that early divergence in maternal gestational weight gain does not yet translate into measurable fetal growth restriction or altered placental scaling.

We next assessed placental features and vascular composition. Quantification of nucleated red blood cells (nRBCs) in the labyrinth layer revealed a significant increase specifically in Fen placentas (Fig. 3E,F), suggesting early fetal or placental stress despite preserved growth metrics. In parallel, labyrinth compartment composition was preserved: the proportions of maternal blood spaces and trophoblast tissue were unchanged between groups (Fig. 3G–H), indicating that gross labyrinth architecture remains intact at this stage.

Because maternal immune–vascular interactions can shape labyrinth perfusion and spiral artery remodeling, we quantified uterine NK (uNK) cells and trophoblast invasion in the decidua. uNK cell abundance was increased in the Fen group (Fig. 3I,J), pointing to altered maternal immune signaling at the implantation site. However, invasive trophoblast abundance did not differ across groups (Fig. 3K), indicating that trophoblast invasion is preserved despite immune alterations. Together, these findings indicate that fentanyl exposure elicits early immune and stress signals at the maternal–fetal interface before detectable placental structural defects or fetal growth restriction emerge.

Consistent with this, spiral artery morphometrics (lumen area) showed no clear differences between groups (Fig. 3L).

### Multidrug exposure affects fetal viability and fetal-placental health metrics

Having identified mid-gestational alterations, we next examined placental and fetal outcomes at term (E18.5) (Fig. 4A). Litter size was unchanged across groups (Fig. 4B), indicating that differences in fetal mass were not attributable to variations in litter size. However, dams exposed to FenTHC showed a significant reduction in the proportion of viable fetuses in relation to unexposed and THC-exposed dams, driven by an increase in late resorptions (Fig. 4C), suggesting that multidrug exposure compromises embryo/fetal viability after placentation.

**Figure 4.**
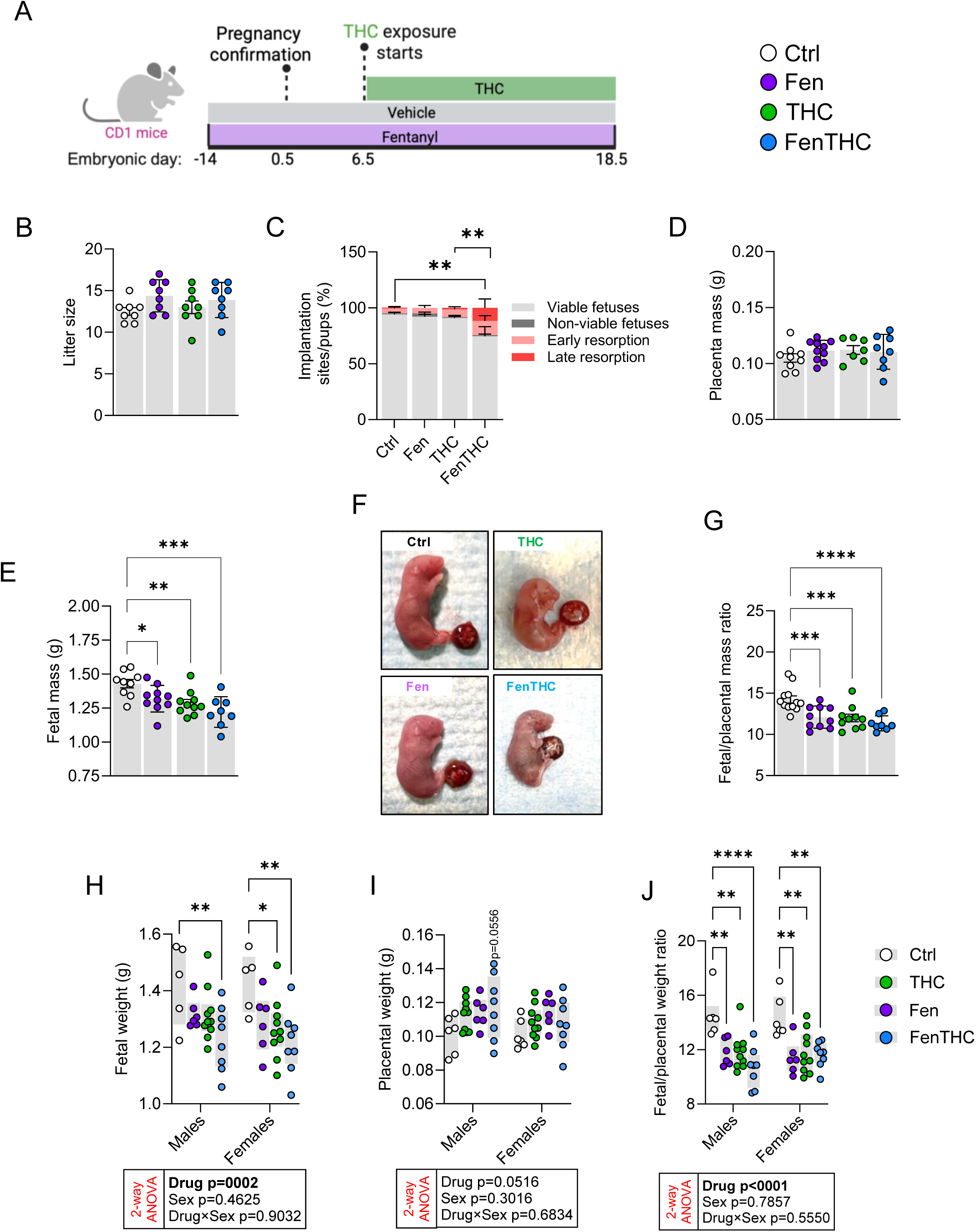
Prenatal fentanyl and Δ^9^-THC exposure reduces fetal growth and placental efficiency at term (E18.5) and combined exposure decreases fetal viability. (A) Experimental design: mice were exposed or not to fentanyl two weeks before pregnancy confirmation at E0.5, fentanyl was administered continuously from E0.5 to E18.5, and THC exposure began at E6.5 and continued through E18.5, with vehicle-treated mice serving as controls. (B) Litter size. (C) Distribution of implantation outcomes, including viable fetuses, non-viable fetuses, early resorptions, and late resorptions. (D) Placental weight at E18.5. (E) Fetal weight. (F) Representative images of fetuses and placentas from each exposure group. (G) Placental efficiency (fetal/placental weight ratio). (H–J) Sex-stratified fetal outcomes, including fetal weight (H), placental weight (I), and placental efficiency (J). Points represent litter means (one point per dam; two points per dam in sex-stratified analyses), and bars show mean ± SEM. Data were analyzed using two-way ANOVA followed by uncorrected Fisher’s LSD post hoc test (A-E, Fen vs THC interaction tests) or Dunnett’s post hoc test comparing treatment groups with controls (G-I, drug exposure vs fetal sex). Ctrl, control; THC, Δ^9^-tetrahydrocannabinol; Fen, fentanyl; FenTHC, fentanyl + Δ^9^-tetrahydrocannabinol. *: p < 0.05, **: p<0.01, ***: p < 0.001, ****: p < 0.0001.

Placental weight was unchanged across groups (Fig. 4D). In contrast, fetal weight was significantly reduced in all drug-exposed groups, with the most pronounced reduction (∼13%) observed in the FenTHC group (Fig. 4E,F). Consequently, placental efficiency (fetal/placental weight ratio) was significantly decreased across all exposure conditions (Fig. 4G), indicating impaired placental support of fetal growth, particularly under multidrug exposure.

No significant main effect of sex was detected for fetal weight, placental weight, or placental efficiency, and no significant drug×sex interactions were observed for any outcome. Drug exposure significantly affected fetal weight; post-hoc comparisons showed lower fetal weight in the FenTHC group in males and females, and in the THC group in females, relative to sex-matched controls (Fig. 4H). Placental weight was not significantly altered by exposure (Fig. 4I). In contrast, placental efficiency (fetal/placental weight ratio) was significantly reduced by drug exposure, with lower ratios in the THC, Fen, and FenTHC groups than in controls in both sexes (Fig. 4J). Overall, these term data indicate that prenatal drug exposure reduced placental efficiency across all exposed groups, whereas lower fetal weight was detected only in specific exposure groups, without evidence of a sex-specific effect.

### Placenta structure is altered in opioids and THC-exposed pregnancies

To determine whether the deficits in fetal growth and placental efficiency were accompanied by structural changes in the placenta, we quantified the major placental compartments at E18.5 (Fig. 5). No significant sex differences were detected in total placental area, decidual area, junctional zone area, or labyrinth area, and no drug×sex interactions were observed for any of these measures (Fig. 5F–I). Drug exposure significantly affected total placental area, decidual area, and labyrinth area, but not junctional zone area. In the pooled analysis, total placental area was increased in the THC and FenTHC groups relative to controls (Fig. 5B), decidual area was increased in the FenTHC group relative to controls and THC (Fig. 5C), and labyrinth area was increased in the THC and FenTHC groups relative to controls (Fig. 5E), whereas junctional zone area was unchanged (Fig. 5D). After stratification by sex, total placental area remained increased in the THC and FenTHC groups in both males and females (Fig. 5F), decidual area remained increased only in the FenTHC group in both sexes (Fig. 5G), and labyrinth area was increased only in the THC group in both sexes (Fig. 5I), while the FenTHC-associated increase seen in the pooled analysis was not retained after sex stratification. Together, these data indicate compartment-specific placental structural changes at E18.5, seen primarily in THC-containing exposure groups, without evidence of sex-dependent effects.

**Figure 5.**
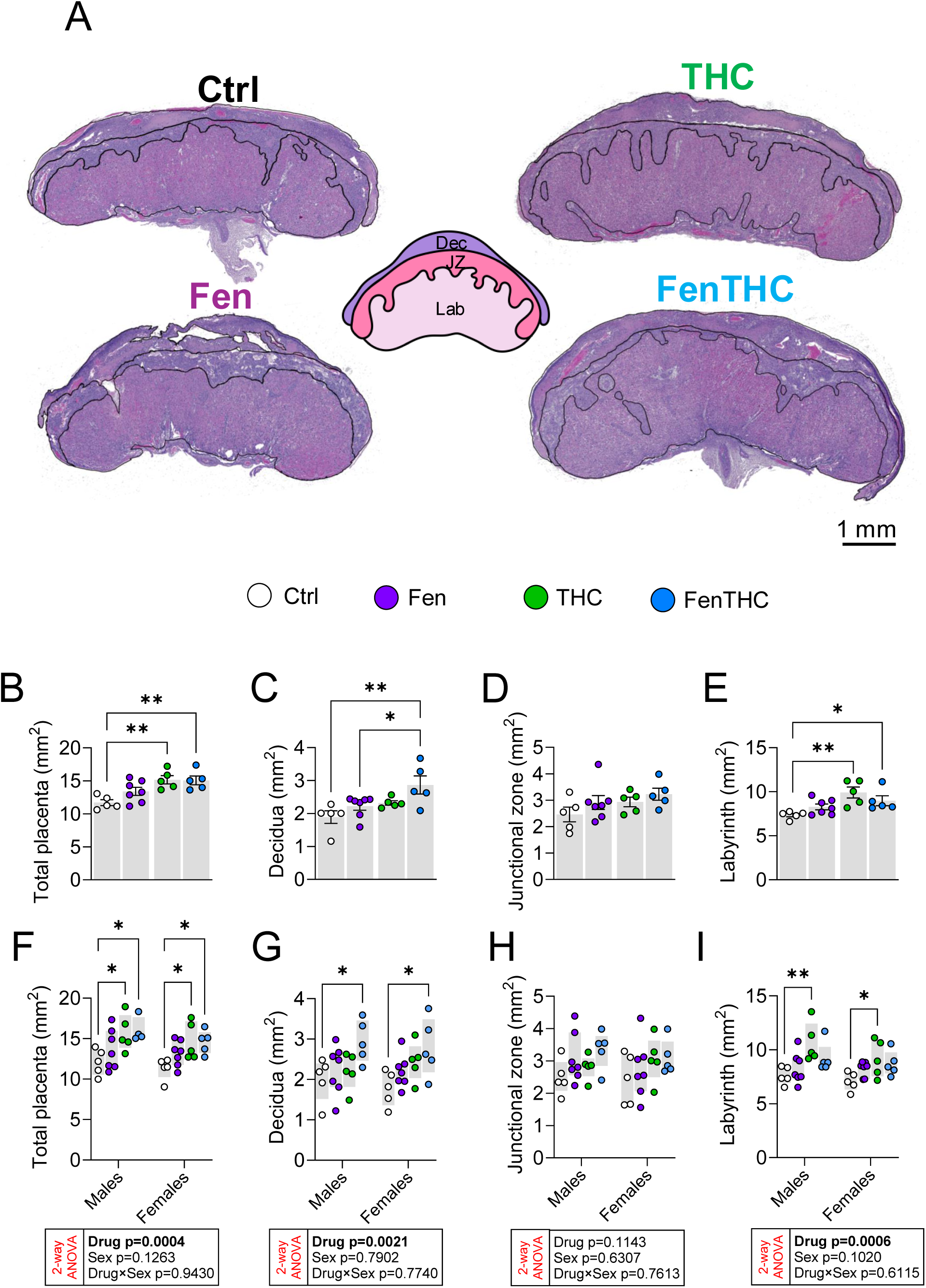
Term placental architecture is remodeled by fentanyl and Δ^9^-THC exposure in a compartment-specific manner. (A) Representative H&E-stained placental cross-sections at E18.5 from each exposure group, with major compartments indicated: decidua (Dec), junctional zone (JZ), and labyrinth (Lab). (B–E) Quantification of total placental cross-sectional area (B) and compartment areas for decidua (C), junctional zone (D), and labyrinth (E). (F–I) Sex-stratified compartment area analyses for total placental area (F), decidua (G), junctional zone (H), and labyrinth (I). Points represent litter means (one point per dam, n=5-7dams/group; two points per dam in sex-stratified analyses), and bars show mean ± SEM. Data were analyzed using two-way ANOVA followed by uncorrected Fisher’s LSD post hoc test (B-E, Fen vs THC interaction tests) or Dunnett’s post hoc test comparing treatment groups with controls (F-I, drug exposure vs fetal sex). Ctrl, control; THC, Δ^9^-tetrahydrocannabinol; Fen, fentanyl; FenTHC, fentanyl + Δ^9^-tetrahydrocannabinol. *: p < 0.05, **: p<0.01.

### Prenatal drug exposure alters placental labyrinth organization and signaling

To determine whether prenatal drug exposure altered the structural and cellular composition of the placental labyrinth, we first examined CD31-stained sections and applied pixel classification to distinguish fetal capillaries (FC), maternal blood spaces (MBS), and tissue (Fig. 6A). Quantitative analysis revealed a significant shift in labyrinth composition in the FenTHC group, characterized by altered proportions of FC, MBS, and tissue area relative to controls, specifically a reduction in the proportion of fetal capillaries and an increase in surrounding tissue coverage (Fig. 6B,C). Ultrastructural examination of the labyrinth by electron microscopy revealed disruption of the feto-maternal interface following drug exposure (Fig. 6D). In control placentas, the trophoblast layer separating fetal capillaries (FC) from maternal blood spaces (MBS) appeared thin, continuous, and well organized, with open and clearly defined vascular channels. In THC- and fentanyl-exposed placentas, this barrier showed mild thickening accompanied by early irregularity and focal shredding of the maternal-facing trophoblast surface into the maternal blood space. These features were markedly exacerbated in the FenTHC group, where pronounced thickening, extensive trophoblast disorganization, and prominent shredding and budding into the maternal compartment were evident. This was associated with apparent swelling, vacuolization, and loss of cellular organization within trophoblast layers, resulting in a structurally disrupted and thick exchange interface between maternal and fetal compartments.

**Figure 6.**
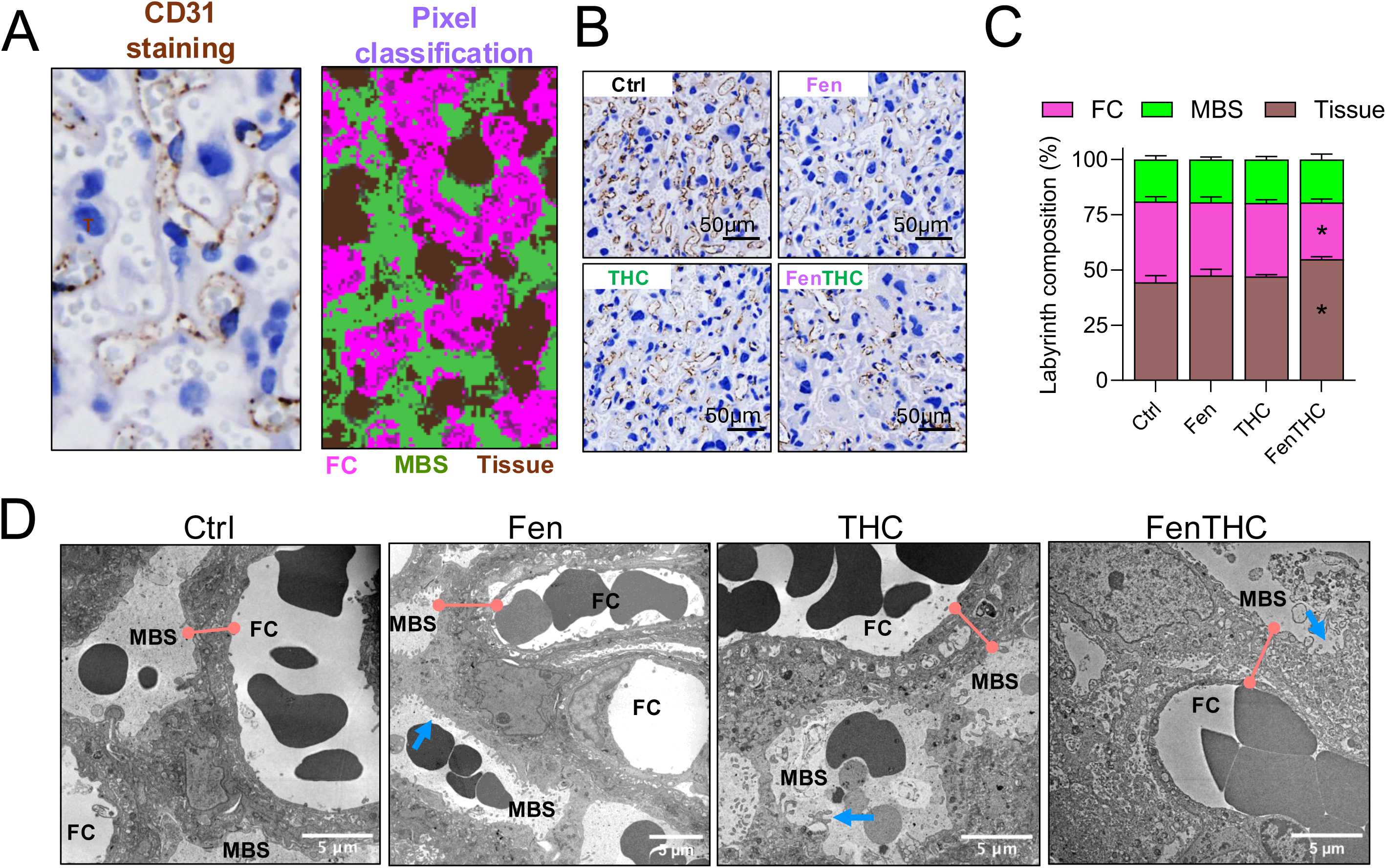
Combined fentanyl+Δ^9^-THC exposure disrupts labyrinth composition and feto-maternal interface ultrastructure at term. (A) Representative CD31 immunostaining and pixel-classification workflow used to segment fetal capillaries (FC; CD31⁺), maternal blood spaces (MBS; CD31⁻ luminal spaces), and trophoblast tissue. (B) Representative CD31-stained labyrinth images from each exposure group. (C) Quantification of labyrinth composition (FC, MBS, tissue) using a pixel classification approach. (D) Representative transmission electron microscopy (TEM) images of the labyrinth showing the barrier between fetal capillaries (FC) and maternal blood spaces (MBS). Blue arrows indicate trophoblast surface irregularities/budding into the maternal blood space, while red lines indicate interhemal barrier thickness. Points represent individual placentas and bars show mean ± SEM. Data were analyzed using two-way ANOVA followed by Dunnett’s post hoc test comparing treatment groups to controls. Placental compartment, fetal sex, and drug exposure were included as factors depending on the analysis (n= 5 dams/group). Interaction terms were tested and are reported in the panels. Ctrl, control; THC, Δ^9^-tetrahydrocannabinol; Fen, fentanyl; FenTHC, fentanyl + Δ^9^-tetrahydrocannabinol. *: p < 0.05, **: p<0.01, ***: p<0.001.

We next examined markers of cellular proliferation, immune signaling, and placental factors relevant to fetal development in placenta samples. Ki67⁺ cell density was significantly affected by drug exposure but not sex (Fig. 7A), indicating decreased proliferative activity across treatment groups. CD68⁺ cell density in the decidua, used as a proxy for placental macrophage abundance, was not significantly affected by drug treatment, fetal sex, or the drug-by-sex interaction. Placental levels of the anti-inflammatory cytokine IL-10 and the pro-inflammatory cytokine IFN-β were significantly influenced by drug treatment (Fig. 7C,D), with reductions that were pronounced following combined drug exposure, consistent with suppression of cytokine-mediated immune signaling within the placenta. Despite these structural and immune alterations, folate concentrations (Fig. 7E) and serotonin (5-HT) levels (Fig. 7F) were unchanged across treatment groups.

**Figure 7.**
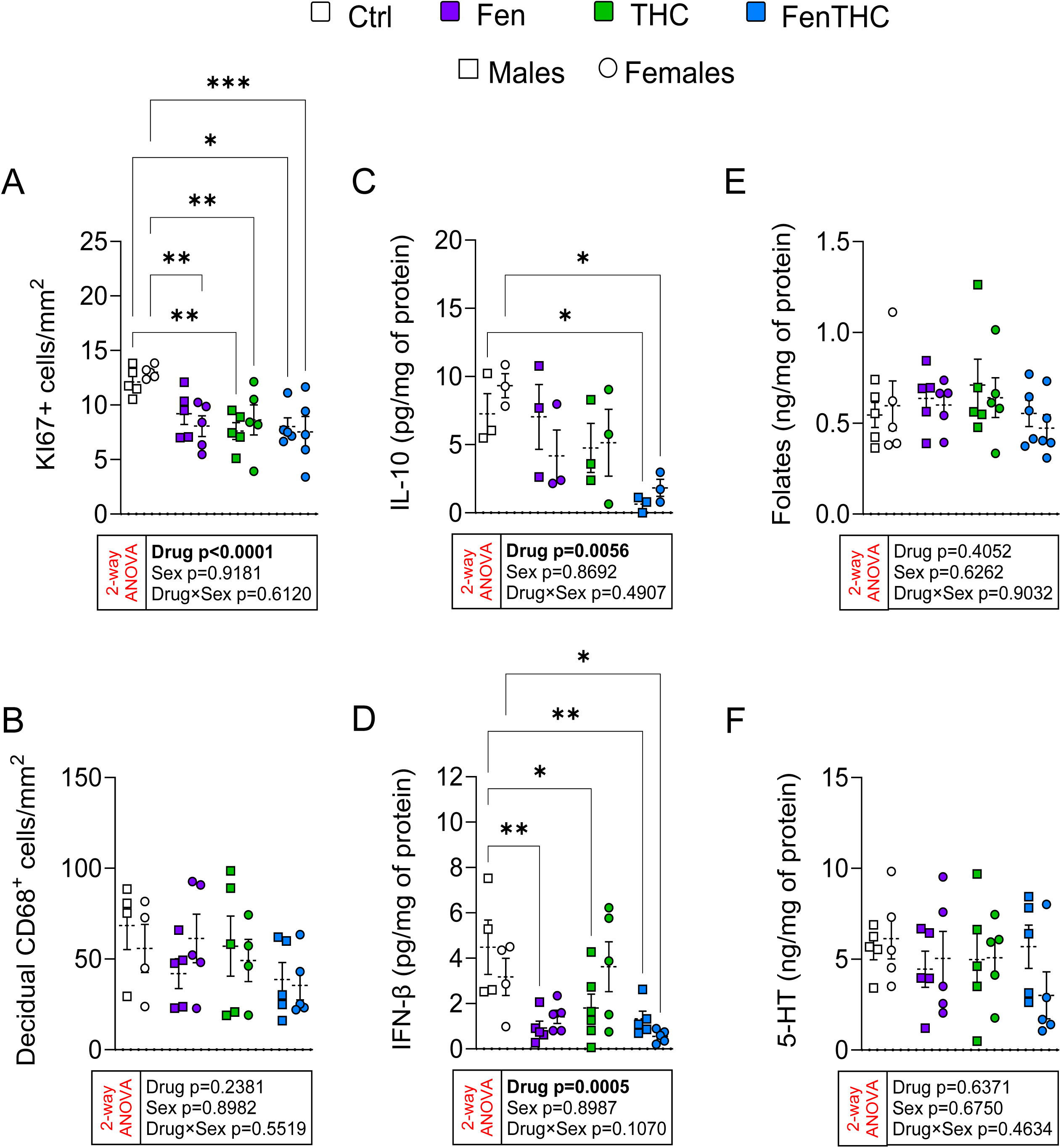
Fentanyl and Δ^9^-THC suppress placental proliferation and cytokine production. (A) Proliferation index quantified as Ki67⁺ cells/mm². (B) Decidual macrophage abundance quantified as CD68⁺ cells/mm². (C-D) Placental cytokine levels measured by ELISA, including IL-10 (C) and IFN-β (D), normalized to total protein. (E-F) Placental folic acid (E) and serotonin (5-HT) (F) levels, normalized to total protein. Points represent individual placentas and bars show mean ± SEM. Data were analyzed using two-way ANOVA followed by Dunnett’s post hoc test comparing treatment groups with controls. Fetal sex and drug exposure were included as factors (n= 3-5 dams/group). Interaction terms were tested and are reported in the panels. Ctrl, control; THC, Δ^9^-tetrahydrocannabinol; Fen, fentanyl; FenTHC, fentanyl + Δ^9^-tetrahydrocannabinol. *: p < 0.05, **: p<0.01, ***: p<0.001.

### Single and multidrug exposure during pregnancy impact placenta transcriptome

Principal component analysis (PCA) was performed on normalized gene expression values to explore global transcriptional variation across treatment groups. We statistically tested the association of each principal component with the experimental variables group and sex. The dominant sources of variance (PC1 and PC2) were not associated with drug exposure (Supplementary Fig. 2). In contrast, drug exposure was strongly associated with PC3, PC4, and PC5 (P adj < 0.01), indicating that treatment effects are reflected in subtler but biologically meaningful transcriptional patterns.

Visualization of PC5 versus PC4, the two components most significantly associated with treatment group, demonstrated clear separation of Ctrl, THC, Fen, and FenTHC samples, with minimal overlap between groups (Fig. 8A). To quantify the magnitude of transcriptomic deviation induced by each treatment, we calculated the Euclidean distance of each sample to the Ctrl group centroid within this PCA space (PC5 vs PC4). Both THC and FenTHC treatments significantly increased transcriptomic distance from Ctrl (Fig. 8B). Sex did not show strong associations with the principal components linked to treatment effects. Instead, sex-related variation appeared to be most represented in PC9. However, PC9 did not exhibit significant differences between treatment groups (Fig. 8C). These results suggest that, although sex contributes to a portion of the overall transcriptional variance, it is not a major driver of the transcriptional patterns associated with prenatal drug exposure.

**Figure 8.**
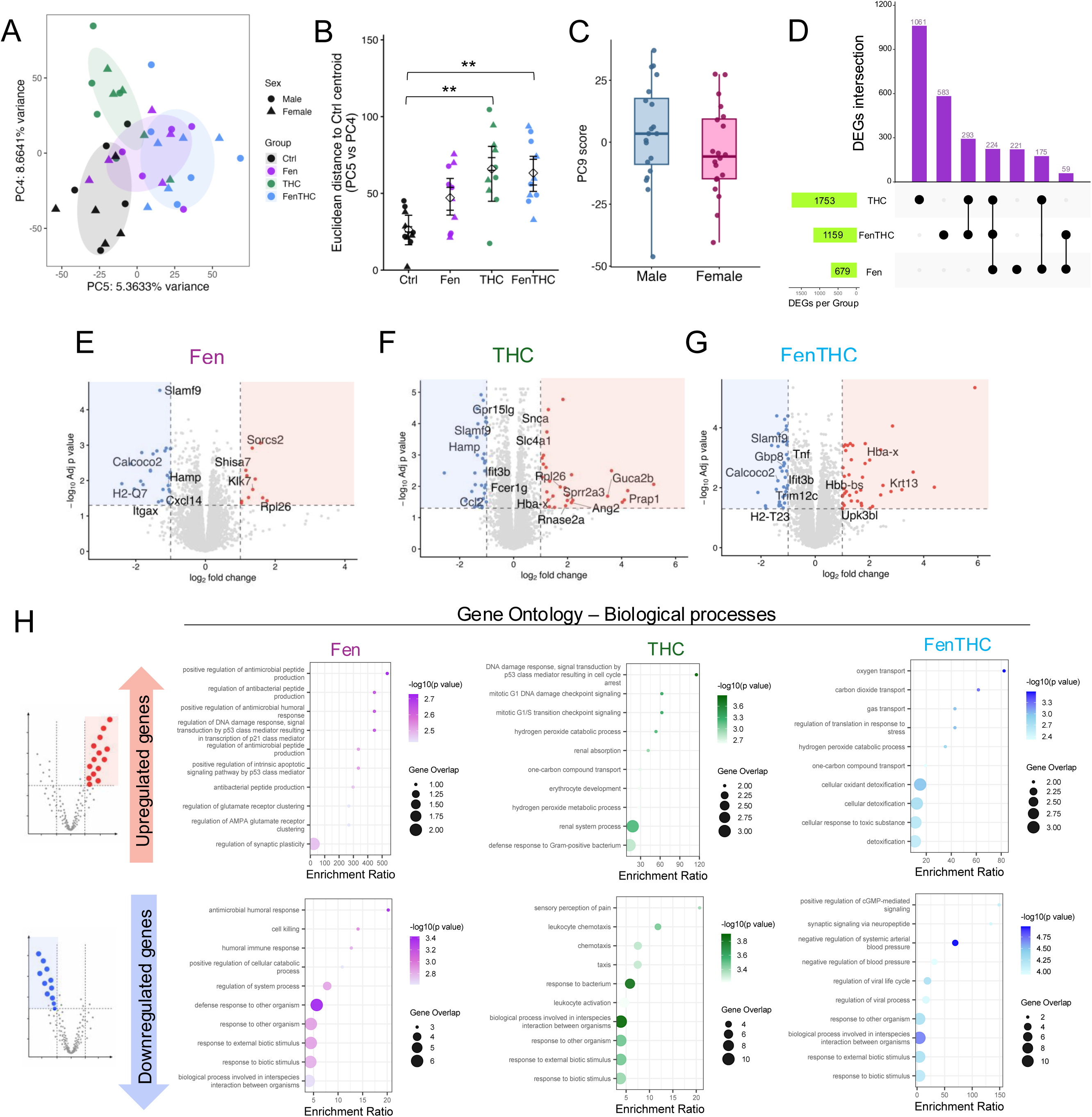
Single and combined drug exposure during pregnancy drives distinct placental transcriptomic programs with limited overlap. (A) Principal component analysis (PCA) of normalized placental gene expression at E18.5 (PC5 vs PC4). Points are colored by exposure group and shaped by fetal sex. (B) Euclidean distance of each sample to the control centroid in PCA space (PC5 vs PC4), quantifying transcriptomic deviation induced by each exposure. (C) PC9 scores stratified by fetal sex. (D) UpSet plot summarizing the number of differentially expressed (DE) genes (P adj < 0.05, green bars) unique to or shared (top, purple bars) between THC, fentanyl, and FenTHC exposures. (E–G) Volcano plots for THC (E), fentanyl (F), and FenTHC (G) contrasts versus control using a stringent threshold (P adj < 0.05 and |log₂FC| > 1). Upregulated (red) and downregulated (blue) genes are highlighted. (H) Gene Ontology (GO) biological process enrichment of upregulated and downregulated DEGs for each exposure group. RNA-sequencing analysis included 10 placenta samples per group (5 male, 5 female). Differential expression was performed with an edgeR-based model including fetal sex as a covariate. The enrichment ratio represents the degree of overrepresentation of genes from each GO pathway among the differentially expressed genes. Dot size indicates gene overlap, and color represents adjusted p-values. For A-C, points represent individual placental samples mean ± SEM. Sal, saline; THC, Δ^9^-tetrahydrocannabinol; Fen, fentanyl; FenTHC, fentanyl + Δ^9^-tetrahydrocannabinol. **p < 0.01, **: p<0.01.

To determine the extent of overlap in treatment-induced transcriptional responses, we performed differential expression analysis –and downstream analyses– for each drug exposure relative to Ctrl while including fetal sex as a covariate to account for residual sex-associated variation. The sets of differentially expressed genes (DEGs) identified for THC, Fen, and FenTHC were then compared using an UpSet intersection analysis (Fig. 8D). For this comparison, DEGs were defined using an adjusted significance threshold of P Adj < 0.05 without applying a fold-change cutoff, allowing visualization of the complete set of genes statistically affected by each treatment. Using this criterion, THC produced the largest transcriptional response (1,753 DEGs), followed by FenTHC (1,159 DEGs) and Fen (679 DEGs). Intersection analysis revealed that most DEGs were treatment-specific, indicating that each drug induces a largely distinct transcriptional program. The largest treatment-specific set consisted of 1,061 genes unique to THC, while FenTHC and Fen showed smaller sets of unique genes: 583 and 221, respectively. Shared transcriptional responses were comparatively limited: 293 genes overlapped between THC and FenTHC, 59 between FenTHC and Fen, 175 between THC and Fen, and only 224 genes were common to all three treatments (Fig. 8D). The interaction analysis (treatment × sex) did not identify any statistically significant interaction effects (Supplementary Fig. 3), indicating that the transcriptional response to exposure was comparable between sexes and showed no evidence of sex-dependent differences. These results indicate that although a modest core response exists across exposures, most transcriptional changes are treatment-specific, with THC driving the most extensive and distinct alterations.

For visualization using volcano plots (Fig. 8E-G) and for functional enrichment analyses (Gene Ontology biological process overrepresentation) (Fig. 8H), we applied a more stringent biological relevance threshold (P Adj < 0.05 and |log₂FC| > 1) to focus on genes exhibiting robust transcriptional changes. THC exposure resulted in coordinated downregulation of genes involved in innate immune activation, leukocyte chemotaxis, and host defense, including key regulators of chemokine signaling (*Ccl2*, *Gpr15lg*), myeloid activation (*Fcer1g*, *Slamf9*), interferon response (*Ifit3b*), and antimicrobial iron regulation (*Hamp*) (Fig. 8E,H). Concurrently, THC induced a transcriptional program consistent with oxidative and genotoxic stress. Enrichment analysis of upregulated genes revealed activation of p53-mediated DNA damage checkpoint pathways (*Rpl26*, *Prap1*), hydrogen peroxide metabolic processes (*Snca*, *Hba-x*), and erythrocyte/oxygen handling genes (*Slc4a1*, *Hba-x*) (Fig. 8F,H). Additional enrichment of renal solute transport and epithelial defense pathways (*Ang2*, *Guca2b*, *Rnase2a*, *Sprr2a3*) indicates disruption of metabolic and barrier homeostasis (Fig. 8F,H). Together, these results indicate that THC exposure may trigger a redox stress response leading to p53-dependent cell cycle arrest and epithelial stress activation in the placenta.

Downregulated genes in fentanyl-exposed placentas were highly enriched for innate immune and antimicrobial defense pathways. This suppression was driven by decreased expression of genes involved in antigen presentation (*H2-Q7*), dendritic/macrophage function (*Itgax*, *Slamf9*), chemokine signaling (*Cxcl14*), antimicrobial iron regulation (*Hamp*), and xenophagy/autophagy-mediated pathogen clearance (*Calcoco2*), indicating impaired placental innate immune defense (Fig. 8 F,H). In contrast, Upregulated genes were enriched for pathways related to antimicrobial peptide regulation (*Klk7*), p53-mediated DNA damage response (*Rpl26*), and glutamate receptor clustering (*Shisa7*, *Sorcs2*), reflecting epithelial stress responses, enhanced p53 translation, and modulation of glutamatergic signaling (Fig. 8F,H). Collectively, these findings indicate that fentanyl exposure induces a coordinated placental stress program characterized by epithelial defense activation, p53-mediated stress signaling, and remodeling of membrane signaling pathways, some components of which are similarly altered in THC-exposed placentas.

FenTHC exposure produced a transcriptional profile characterized by suppression of placental innate immune surveillance and host–pathogen response pathways. Downregulated genes were strongly enriched for biological processes related to host defense against pathogens and innate immune responses (Fig. 8G,H). These enrichments were driven by coordinated downregulation of genes involved in myeloid immune signaling (*Slamf9*, *Tnf*), interferon-stimulated and antiviral defense responses (*Ifit3b*, *Gbp8*, *Trim12c*, *H2-T23*), and xenophagy/autophagy-mediated pathogen clearance (*Calcoco2*). In contrast, genes upregulated in FenTHC-exposed placentas were enriched for oxygen transport, oxidant detoxification, hydrogen peroxide catabolism, and cellular responses to toxic substances (Fig. 8G,H), driven by increased expression of genes associated with oxygen handling (*Hba-x*, *Hbb-bs*), oxidative stress responses (*Upk3bl*), and epithelial remodeling (*Krt13*). Together, these results indicate that FenTHC exposure suppresses placental innate immune defense programs while simultaneously promoting transcriptional responses linked to oxidative stress, detoxification, and cellular remodeling.

Across treatments, THC and fentanyl—both alone and in combination—consistently suppressed placental innate immune and antimicrobial defense programs, indicating a shared reduction in immune surveillance in response to drug exposure. This transcriptional pattern aligns with the observed drug-associated decreases in placental IL-10 and IFN-β levels, supporting an overall attenuation of cytokine-mediated immune activity within the placenta.

### Prenatal drug exposure induces shared immune suppression and treatment-specific cellular remodeling programs

Gene Set Enrichment Analysis (GSEA) performed on the full ranked transcriptome, for detection of subtle but biologically meaningful shifts in gene programs, confirmed that all treatments induced a coordinated suppression of innate immune and host-defense pathways, including interferon response, cytokine signaling, and response to bacteria and viruses (Fig. 9A-C). Despite this shared suppression, the transcriptional programs activated by each treatment were distinct. THC preferentially enriched pathways related to RNA processing, mRNA metabolism, and chromosome organization, consistent with activation of nuclear stress and transcriptional regulation mechanisms (Fig. 9A). Fen exposure enriched pathways associated with ribosome biogenesis, RNA transport, and translation, indicating activation of protein synthesis machinery (Fig. 9B). In contrast, FenTHC exposure strongly enriched pathways related to cell projection organization, vesicle trafficking, glycosylation, and metabolic transport, reflecting structural and metabolic remodeling of placental cells (Fig. 9C). These findings demonstrate that while drug exposure consistently suppresses placental immune surveillance, each treatment elicits a distinct adaptive cellular response.

**Figure 9.**
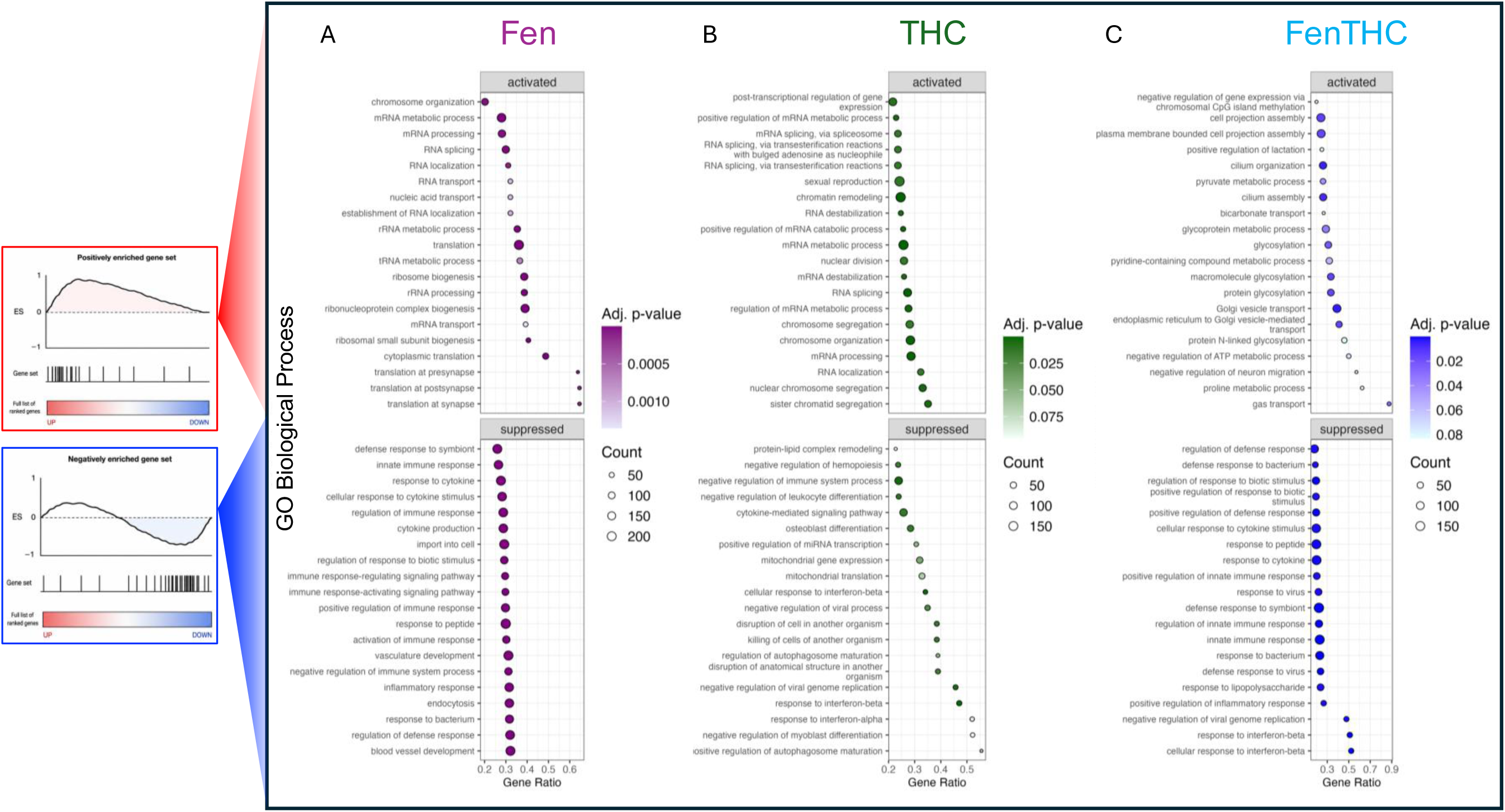
Gene set enrichment analysis reveals shared immune suppression and treatment-specific stress/metabolic responses. Gene set enrichment analysis (GSEA) of the full ranked placental transcriptome for GO Biological Process pathways. (A) GSEA dot plot for the THC versus control contrast. (B) GSEA dot plot for the fentanyl versus control contrast. (C) GSEA dot plot for the fentanyl+THC versus control contrast. Dot size represents the number of genes contributing to each pathway (Count), and color indicates the adjusted p-value. Pathways are separated into those enriched as positively versus negatively enriched and plotted according to their gene ratio, which reflects how enriched the pathway is relative to the gene list size. Sal, saline; THC, Δ^9^-tetrahydrocannabinol; Fen, fentanyl; FenTHC, fentanyl + Δ^9^-tetrahydrocannabinol.

### Fen×THC interaction reveals synergistic lipid responses and lower-than-additive RNA-processing programs in placenta

To determine whether adding THC to chronic fentanyl exposure produces a placental transcriptional response different from that predicted by the individual exposures, we performed an edgeR interaction analysis using the contrast (FenTHC − Fen) − (THC − Ctrl) while adjusting for fetal sex. This identified 933 genes with significant interaction effects (P adj < 0.05; 423 positive and 510 negative coefficients) (Fig. 10A). Ranked GSEA across all interaction coefficients showed that positive interaction signals were enriched for glycerophospholipid/glycerolipid metabolism and phosphatidylcholine catabolism, whereas negative interaction signals were enriched for RNA splicing, mRNA processing, ribonucleoprotein complex biogenesis, rRNA metabolism, and ribosome biogenesis (Fig. 10B). A more stringent significance cutoff (P adj < 0.05 and |log₂FC| > 1) revealed 63 genes with interaction effects used for visualization (Fig. 10D). Positive interaction coefficients indicate genes whose expression in the combined exposure was higher than expected under the additive model, consistent with synergistic co-exposure effects, whereas negative interaction coefficients indicate genes whose expression was lower than expected under additivity. Representative positive interaction genes included *Rpl21*, *Gdpd3*, *Trim34a*, *Smpd1*, and *Slamf9*, while negative interaction genes included *Trim68*, *Cebpb*, *Trim12c*, *Snca*, *Rpl26*, and *Gcm1*. Consistent with this, the GO overrepresentation analysis of positive interaction genes (right panel) suggested enrichment of lipid- and membrane-related processes, whereas the analysis of negative interaction genes (left panel) highlighted broader cellular organization and catabolic terms.

**Figure 10.**
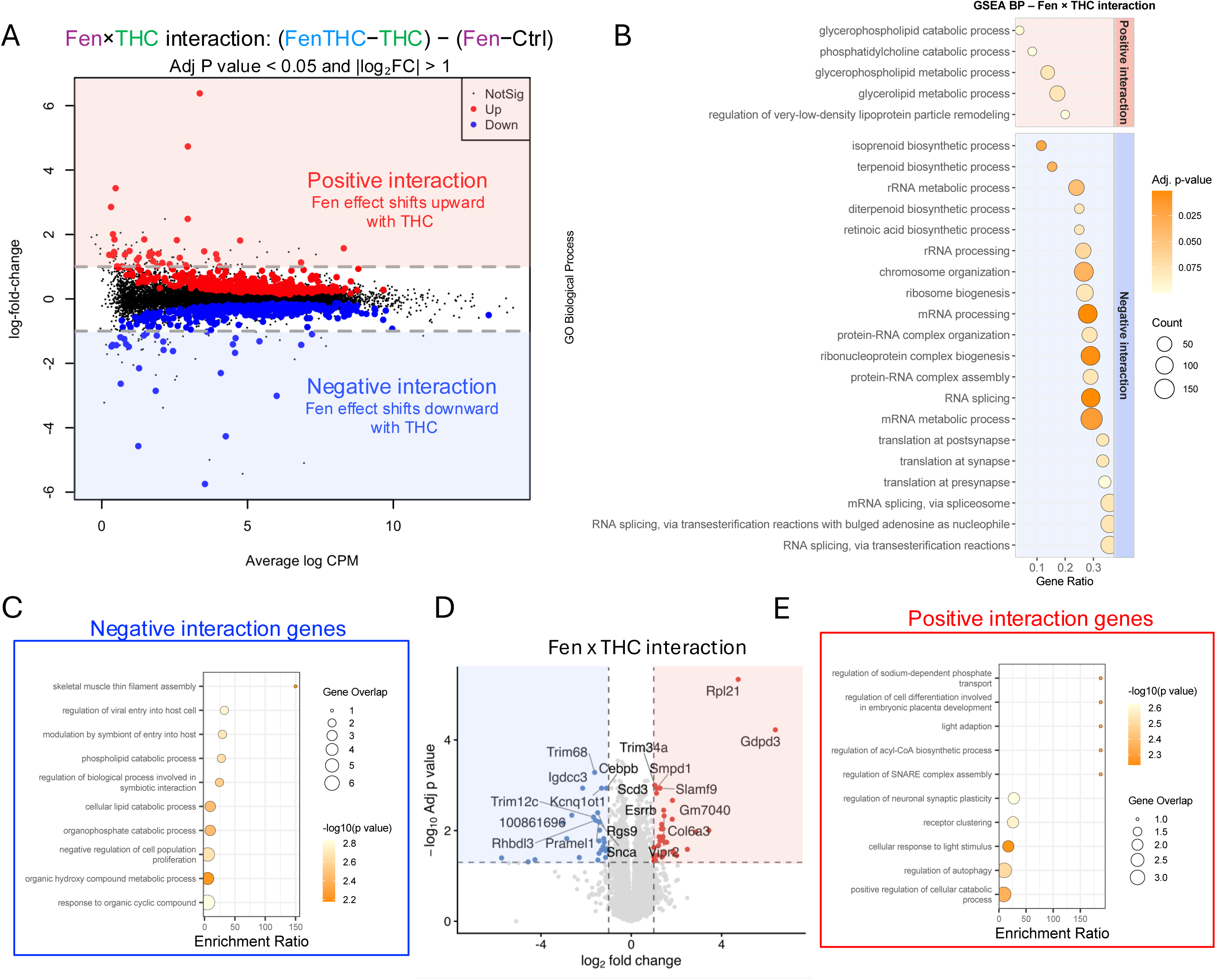
Combined opioid–cannabinoid exposure drives interaction-specific transcriptomic reprogramming. (A) Scatter plot showing the Fen×THC interaction contrast (FenTHC − Fen) − (THC − Saline), with log_2_ fold change plotted against average log counts per million (CPM). Genes are colored as upregulated (red), downregulated (blue), or not significant (black). Dashed lines indicate significance thresholds. (B) Gene set enrichment analysis (GSEA) of biological processes associated with the Fen×THC interaction. Dot size reflects gene count within each pathway, and color represents adjusted p-values. Gene ratio indicates the proportion of differentially expressed genes within each pathway. (C,E) Gene Ontology (GO) enrichment analysis of negatively (C) and positively (E) interacting genes (. Dot size indicates gene overlap, and color corresponds to −log_10_(p-value). (D) Volcano plot highlighting key differentially expressed genes in the interaction contrast. Selected genes of interest are labeled. Vertical and horizontal dashed lines indicate fold change (|log2FC| = 1) and significance thresholds (P adj <0.05), respectively.

The enrichment of *Gdpd3* and *Smpd1* supports altered phospholipid and sphingomyelin turnover under combined exposure, providing a potential molecular link to the trophoblast membrane budding and disruption of the maternal–fetal exchange interface observed by electron microscopy. The negative interaction involving *Gcm1* is also notable because *Gcm1* regulates syncytiotrophoblast differentiation and labyrinth branching. Together, these findings indicate that adding THC to chronic fentanyl exposure alters placental membrane-lipid metabolism and RNA-processing programs in ways that are not predicted by either exposure alone.

### Combined opioid and cannabinoid exposure most strongly perturbs growth-linked placental gene networks

To identify placental transcriptional programs intrinsically linked to fetal growth in the context of single and multidrug exposure, we performed sex-adjusted partial correlation analysis between normalized gene expression and average fetal weight across all samples (Fig. 11A). The distribution of partial correlations revealed a distinct subset of genes strongly associated with fetal weight (|r| up to ∼0.8, P Adj < 0.05). These genes segregated into two opposing groups: genes positively correlated with fetal weight (higher expression in larger fetuses) and genes negatively correlated with fetal weight (higher expression in growth-restricted fetuses), defining two transcriptional programs intrinsically linked to placental growth capacity. Genes positively associated with fetal weight included regulators of vascular patterning, morphogenesis, and transcriptional control (i.e. *Bmp4*, *Wnt7b*, *Gata2*, *Nrp1*, *Eng*, *Hoxa1*, *Mdk*, *Igfbp6*), whereas genes negatively associated with fetal weight were enriched for hypoxia/erythroid signatures and metabolic stress responses (i.e. *Vegfa*, *Hba-x*, *Gdpd3*, *Prl2b1*, *Bcl2l2*, *Vps39*). This organization suggests that placentas associated with fetal growth restriction under cannabis and opioid exposure shift toward a compensatory stress-adaptation state, with potential vascular compromise, that correlates with reduced fetal weight.

**Figure 11.**
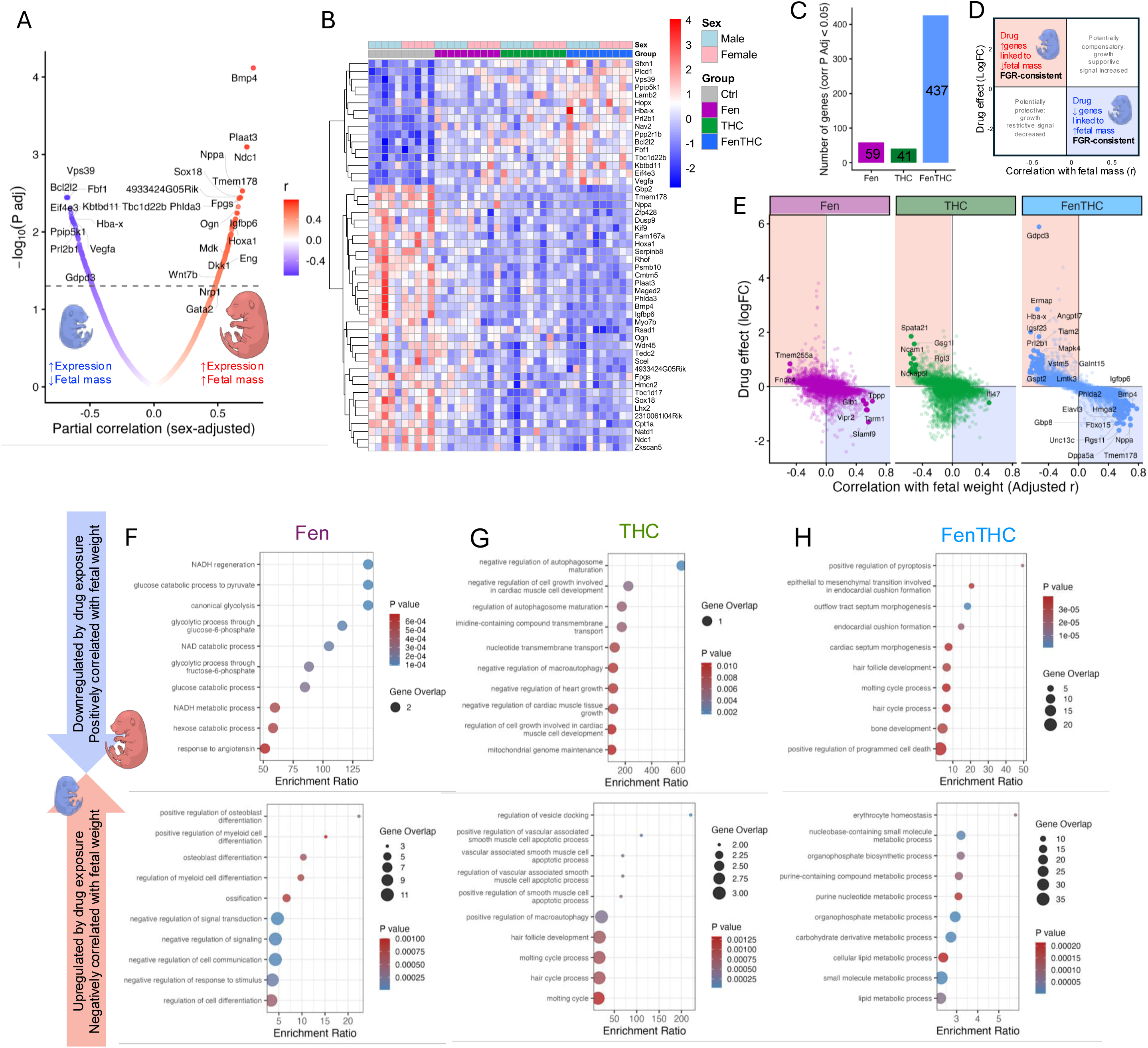
Combined fentanyl+Δ^9^-THC exposure most strongly perturbs fetal-growth–linked placental gene networks. (A) Sex-adjusted partial correlation analysis between placental gene expression and average fetal weight across all samples, showing correlation coefficients (x-axis) and significance (y-axis, adjusted p values). (B) Heatmap of the top 50 fetal weight–associated genes (based on adjusted p values), showing z-scored expression across samples grouped by exposure and fetal sex. (C) Integration of fetal weight correlations with treatment-induced differential expression. Scatter plot shows each fetal-weight–associated gene plotted by its correlation with fetal weight (x-axis) and the log_2_ fold change of its primary driver exposure (y-axis; driver defined as the exposure with the largest absolute log_2_FC among THC, fentanyl, and fentanyl+THC contrasts). Dashed reference lines (|r| = 0.3 and |log2FC| = 0.5) were included as visual guides to highlight genes with moderate correlation strength and biologically meaningful expression changes; these thresholds were not used for statistical filtering. Bar plot summarizes the number of significantly growth-associated genes primarily driven by each exposure. (D–F) GO biological process functional enrichment of fetal weight–associated genes partitioned into growth-supportive genes (positively correlated with fetal weight and downregulated by exposure; top) versus growth-restrictive genes (negatively correlated with fetal weight and upregulated by exposure; bottom) for THC (D), Fen (E), and FenTHC (F). The enrichment ratio represents the degree of overrepresentation of genes from each GO pathway among the differentially expressed genes. Dot size indicates gene overlap, and color represents adjusted p-values. Ctrl, control; THC, Δ^9^-tetrahydrocannabinol; Fen, fentanyl; FenTHC, fentanyl + Δ^9^-tetrahydrocannabinol.

Heatmap visualization of the top fetal weight–associated genes demonstrated a progressive inversion of these programs across treatments (Fig. 11B). Genes supporting vascularization, transcriptional regulation, and growth—such as *Bmp4*, *Wnt7b*, *Gata2*, *Nrp1*, *Eng*, *Hoxa1*, *Igfbp6*, *Fpgs* (critical for folate balance and cell proliferation) and *Ogn* (a growth promoting factor)—were higher in control placentas and progressively suppressed in THC, Fen, and most strongly in FenTHC. In contrast, stress- and restriction-associated genes—including *Vegfa*, *Hba-x*, *Gdpd3*, *Prl2b1*, *Bcl2l2*, and *Vps39*—showed the opposite pattern, with increased expression in drug-exposed placentas, particularly in the combined exposure group (Fig. 11B).

To determine which treatment most strongly influenced each growth-associated gene, we integrated the correlation results with differential expression from each treatment contrast. For each gene, the treatment producing the largest absolute log fold change was assigned as the primary driver (Fig. 11C). Among genes significantly correlated with fetal weight, FenTHC most frequently drove expression changes, followed by Fen and then THC (Fig. 11C), indicating that combined exposure most strongly perturbs genes linked to placental growth in this study. Plotting the correlation of each gene with fetal weight against the drug-induced logFC of its primary driver generated a mechanistic map of treatment effects (Fig. 11C). Genes in the upper-left quadrant (negatively correlated with fetal weight and upregulated by drug exposure) and lower-right quadrant (positively correlated with fetal weight and downregulated by drug exposure) represent transcriptional changes consistent with fetal growth restriction or proper fetal growth, respectively, in this mouse model.

Gene ontology enrichment of fetal-weight–associated genes revealed that each treatment shifts the placenta away from growth-supportive programs toward growth-restrictive programs, but through distinct biological routes (Fig. 11D–F). In THC-exposed placentas (Fig. 11D), genes positively correlated with fetal weight that were suppressed by treatment were enriched for autophagosome maturation, membrane and nucleotide transport, and mitochondrial genome maintenance, suggesting reduced expression of trafficking- and mitochondrial homeostasis–related programs associated with higher fetal weight. Conversely, genes negatively correlated with fetal weight and upregulated by THC were enriched for vascular smooth muscle–associated, cytoskeletal, and vascular structural pathways, consistent with increased expression of structural remodeling programs associated with lower fetal weight. In Fen-exposed placentas (Fig. 11E), genes suppressed by treatment were strongly enriched for NADH regeneration, glycolysis, glucose and hexose metabolism, and broader energy-producing metabolic pathways. In contrast, genes upregulated by Fen were enriched for cell differentiation, osteoblast/myeloid differentiation, and regulation of signaling and cell communication, suggesting a shift away from energy metabolism toward differentiation- and signaling-related programs. FenTHC exposure (Fig. 11F) was associated with the broadest transcriptional reorganization. Genes suppressed by FenTHC were enriched for cell projection organization, epithelial-to-mesenchymal transition–like processes, autophagy, structural morphogenesis, and positive regulation of programmed cell death, consistent with extensive membrane remodeling, structural reorganization, and enrichment of cell death–associated pathways. At the same time, genes upregulated by FenTHC were enriched for purine and nucleotide metabolism, organophosphate and small-molecule metabolism, and lipid metabolic processes, indicating substantial metabolic and biosynthetic reprogramming.

Together, these patterns suggest that all treatments shift placental gene expression away from programs associated with fetal growth. THC was associated primarily with reduced vascular and cytoskeletal organization, Fen with suppression of energy metabolism and increased differentiation- and signaling-related pathways, and FenTHC with broader metabolic reprogramming accompanied by structural remodeling and enrichment of cell death–related processes.

## Discussion

Opioid and cannabis co-use is increasingly common among individuals of reproductive age, yet how these exposures affect placental function and fetal growth remains poorly understood—particularly when drug use precedes pregnancy. Using a mouse model of fentanyl and THC exposure, we show that although pregnancy dampens overt maternal metabolic disturbances, drug exposure produces a pronounced placental phenotype characterized by structural remodeling of placental compartments and the feto-maternal exchange interface, suppression of immune and cytokine signaling, and treatment-specific transcriptional reprogramming. These placental alterations closely track with reduced placental efficiency and fetal growth restriction, identifying the placenta as a primary site where prenatal drug exposure exerts its effects. Notably, combined fentanyl and THC exposure consistently produced the most severe disruption across structural, immune, and molecular measures, indicating that multidrug exposure most strongly reprograms placental function in ways that constrain fetal growth.

Initial metabolic profiling in non-pregnant females demonstrated that combined fentanyl and THC exposure substantially disrupted baseline physiology, reducing respiratory activity, shifting substrate utilization toward fat oxidation, and lowering overall energy expenditure. These findings indicate that multidrug exposure directly alters systemic metabolic regulation outside the context of pregnancy. Strikingly, when the same exposures occurred during gestation, these metabolic disturbances were largely attenuated at E16.5. Pregnant dams exposed to fentanyl displayed only transient hyperlocomotion followed by brief respiratory changes—responses previously reported following fentanyl exposure(40) —with no sustained evidence of metabolic suppression, suggesting that pregnancy buffers or compensates for systemic metabolic perturbations induced by the drugs.

In contrast, the increased locomotion observed during the dark phase in THC-exposed dams is consistent with previously described THC-induced hyperlocomotion at low doses, mediated by cannabinoid receptor 1 (CB1) signaling in striatal neurons that regulate locomotor circuits(41). Alternatively, this pattern may reflect a behavioral response associated with cannabinoid withdrawal or altered arousal state(42). This interpretation is consistent with reports showing altered pharmacokinetics during pregnancy, including increased metabolic clearance of both fentanyl(43) and THC(44), as well as evidence that repeated exposure can lead to physiological tolerance to acute respiratory and behavioral effects(45, 46). While THC consumption is often assumed to increase appetite(47) and fentanyl to reduce it(48), no sustained changes in food intake were observed in the present study. Previous work indicates that these effects are highly dependent on dose and route of administration(47, 49). Despite this apparent buffering of maternal physiology and no changes in food intake, reduced gestational weight gain and fetal growth restriction still emerged. This dissociation between maternal metabolic stability and impaired fetal growth points to the placenta, rather than the maternal metabolic system, as the primary site where drug exposure manifests its pathological effects during gestation.

Evidence of altered placenta composition was already apparent by mid-gestation. At E13.5, increased nRBCs in Fen placentas are consistent with heightened fetal or placental hypoxic stress. In parallel, Fen exposure increased uNK cell abundance at mid-gestation. uNK cells regulate placental vascular remodeling and perfusion, and macrophages contribute to tissue remodeling and inflammatory signaling at the maternal–fetal interface(50–52). Their expansion may reflect a compensatory response to impaired oxygen delivery within the labyrinth, consistent with elevated nRBCs.

Notably, however, this increase in immune cell abundance did not appear to translate into enhanced cytokine signaling, as placental IL-10 and IFN-β levels were reduced later in gestation. Because uNK cells act primarily through cytokine and growth factor secretion(53), this dissociation suggests altered immune cell function that is compensated by increased cell recruitment. This interpretation is supported by recent findings in pregnancies affected by OUD, where increased decidual monocytes and uNK cells were accompanied by reduced NK degranulation and impaired macrophage responses to bacterial stimulation, reflecting disrupted trophoblast–immune communication essential for vascular development(54). Consistent with this, THC alone or in combination with fentanyl did not alter the abundance of the placental immune cell populations quantified in our study, in line with a recent report showing that uNK cell numbers were unaffected by THC but increased by CBD(55). Importantly, in that study, THC-induced fetal growth restriction was associated with reduced production of angiogenic factors such as IFN-γ and VEGF by uNK cells(55). In our model, neither trophoblast invasion nor spiral artery remodeling was altered across drug exposure conditions, indicating that the primary defect is not related to implantation or maternal vascular transformation but rather to the organization and perfusion of the feto-maternal exchange region. These early hypoxia-associated and immune-regulatory changes induced by fentanyl appear to precede the more pronounced structural and transcriptional abnormalities observed at term.

By E18.5, this progressed to a phenotype of growth-restricted fetuses and reduced placental efficiency (56). Placenta functional layers was impacted, with THC and FenTHC selectively expanding the labyrinth—the primary site of feto-maternal exchange, consistent with previous rat studies of gestational THC exposure showing labyrinth enlargement, vascular disorganization, and reduced GLUT1 expression(57)— while THC and FenTHC increased decidual area without affecting the junctional zone, which may reflect delayed decidual regression or altered decidual differentiation programs, consistent with reduced expression of *Bmp4* and *Hoxa1*—both positively associated with fetal mass in our dataset. Such disproportionate placental growth is widely recognized as a compensatory but functionally inadequate response in fetal growth restriction. Notably, FenTHC uniquely enriched gene programs related to cell projection organization, plasma membrane–bounded projection assembly, and vesicle trafficking, transcriptional signatures that are consistent with trophoblast membrane budding and surface shredding observed ultrastructurally by electron microscopy. This pattern could reflect increased vesicle trafficking and projection organization as an adaptive but ultimately destabilizing response of trophoblast cells to drug exposure.

Clinically, these findings align with a model where opioid exposure alone may not uniformly produce gross placental enlargement, but polysubstance co-exposures can worsen placental health. In a prenatal MRI study of pregnancies receiving medication-assisted treatment for opioid use disorder, placental volume did not differ significantly from controls overall; however, concomitant smoking and polysubstance exposure were suggested to be detrimental to placental health(58). Together with our observation that combined Fen and THC produced the most severe structural, immune, and molecular disruption, these data support the idea that co-use patterns may be a key determinant of placental vulnerability and growth restriction risk.

Placental transcriptomic profiling revealed that all drug exposures were characterized by a shared suppression of innate immune and antimicrobial defense programs, including interferon signaling, cytokine production, and responses to bacterial and viral stimuli.

This pattern is consistent with prior reports of immune dysregulation in human placentas following prenatal cannabis exposure(59) and aligns with evidence of impaired immune function associated with both opioid(60, 61) and cannabis(62, 63) use. In our study, this transcriptional signature closely paralleled the observed reductions in placental IL-10 and IFN-β levels, indicating functional suppression of immune signaling at the feto-maternal interface. Such drug-induced immunosuppression may increase susceptibility to infectious diseases during pregnancy. Previous studies have linked cannabis exposure to heightened vulnerability to viral infections, including SARS-CoV-2(64) and bacterial infections such as *Clostridioides difficile*(65). Similarly, opioid use disorder in pregnancy is strongly associated with increased prevalence of hepatitis C virus (HCV) infection at delivery(66), reflecting the high cumulative infection risk among individuals with a history of opioid injection, where nearly half of injecting users—and a substantial proportion of prescription opioid users—acquire HCV within five years of use(67, 68).

Despite the increased abundance of uNK cells and macrophages observed in fentanyl-exposed placentas, the concurrent suppression of cytokine and interferon programs suggests that immune cell recruitment occurs alongside functional impairment. This interpretation is consistent with recent findings in opioid-exposed human pregnancies, where decidual immune cells are numerically increased but exhibit reduced degranulation capacity and impaired macrophage responses to immune stimulation(54).

Consistent with the idea that opioids can target specific trophoblast cel types, oxycodone exposure in pregnant mice has been reported to disrupt invasive parietal trophoblast giant cells (pTGCs)—a lineage with parallels to human EVT biology—and to reduce expression of endocrine/metabolic program genes in that compartment(69). In our model, we did not detect impaired trophoblast invasion or spiral artery remodeling, suggesting the dominant defect lies in exchange-zone organization and function. More broadly, trophoblastic opioid receptor expression and opioid-responsive transcriptional targets can vary with trophoblast differentiation state(70), suggesting cell-type–specific susceptibility of villous trophoblasts to opioid effects and a plausible reason for inter-study variability.

THC and fentanyl, alone and in combination induced broad transcriptional reprogramming in placenta accompanied by suppression of immune signaling pathways, but the underlying stress responses differed. THC exposure produced a signature of oxidative, mitochondrial, and genotoxic stress, marked by increased *Snca*, *Hba-x*, and activation of p53-dependent DNA damage and cell-cycle arrest pathways (*Rpl26*, *Prap1*). Previous studies have found that DEGs in placenta after THC use are related to cytokine binding, regulation of cell migration, cell-substrate adhesion, angiogenesis, and vascular development in non-human primates model(71). In addition, placental induction of fetal hemoglobin genes (*Hba x*/*Hbb y*) under hypoxic challenge has been reported to correlate inversely with fetal mass(72), supporting the interpretation that elevated *Hba x* in our growth-restricted placentas likely reflects a hypoxia or stress-associated programs. In contrast, fentanyl exposure suppressed vascular development programs and activated nucleolar, ribosomal, and translational pathways, indicating a stress response centered on maintenance of protein synthesis.

Combined fentanyl and THC exposure generated a distinct and more severe profile characterized by enrichment of redox homeostasis (*Hba-x*, *Aldh1l1/2*, *Nampt*), glycolysis and β-oxidation (*Gpi1*, *Pkm*, *Pfkp*, *Acadsb*, *Acaa2*), nucleotide turnover (*Upp1*, *Nudt16*, *Pfas*), and lipid remodeling (*Fasn*, *Hmgcr*, *Ppara/Ppard*). These pathways reflect metabolic reprogramming toward energy production and oxidative stress adaptation. Concurrent enrichment of cell projection organization, vesicle trafficking, and membrane remodeling paralleled the trophoblast budding and surface disorganization observed ultrastructurally. Importantly, Fen×THC interaction analysis further indicated that fentanyl does not simply add to the placental effects of THC, but selectively reshapes them by enhancing higher-than-additive lipid-related responses while dampening broader RNA-processing and ribosome-associated programs.

Linking gene expression to fetal growth revealed a reversal of normal placental programs. Genes associated with vascular patterning and developmental signaling (*Bmp4*, *Wnt7b*, *Gata2*, *Nrp1*, *Eng*, *Hoxa1*, *Ogn*, *Fpgs*) were progressively downregulated across treatments, whereas pathways related to hypoxia, metabolic stress, and cellular survival were upregulated. Gene ontology analysis further indicated that these growth-associated transcriptional changes were driven through distinct mechanisms by each exposure: THC was associated with reduced trafficking- and mitochondrial homeostasis–related programs linked to higher fetal weight, together with increased vascular and cytoskeletal remodeling pathways associated with lower fetal weight; fentanyl was associated with suppression of energy metabolism and enrichment of differentiation- and signaling-related pathways; and combined exposure, which showed the broadest set of correlated genes, was associated with structural remodeling, autophagy, programmed cell death–related processes, and broader metabolic reprogramming, including lipid and purine metabolism.

Epigenetic programming may provide an additional mechanism linking prenatal opioid exposure to downstream fetal neurodevelopment. A pilot human study reported opioid-associated placental DNA methylation differences enriched for neurodevelopment/synaptic gene pathways, supporting the concept that placental epigenetic remodeling could contribute to placental dysfunction and neurobehavioral outcomes, including neonatal withdrawal phenotypes(73). At the same time, placenta-mediated mechanisms do not exclude direct fetal neurodevelopmental effects of opioids; for example, methadone has been reported to increase apoptosis in neurons and oligodendrocytes and to promote proinflammatory activation of glial cells in vitro(74). Although folate concentrations were unchanged here, growth-linked expression of folate-pathway genes suggests that one carbon metabolism may still be involved at the level of flux or compartmentalization. Given reports that both deficient and excessively high folic acid exposure can alter offspring neurodevelopment in animal models(75, 76), future work should explicitly consider nutritional context when evaluating drug-associated fetal programming. Together, these studies motivate future work that integrates placental structure/transcriptomics with placental epigenetic profiling and fetal brain endpoints under single vs combined drug exposures.

Several limitations should be considered. Transcriptomic analysis was performed at a single gestational time point and does not capture dynamic changes across pregnancy. Functional measures of placental hypoxia, perfusion and exchange capacity were not directly assessed. As with all murine models, translation to human pregnancy requires caution. Future studies should evaluate longitudinal transcriptional changes, functional exchange measurements, and validation in human placental tissues.

Together, these findings demonstrate that prenatal opioid and cannabis exposure does not primarily disrupt maternal systemic physiology during pregnancy but instead reprograms the placenta at structural, immune, and transcriptional levels. This reprogramming produces a growth-restrictive placental state that is most pronounced under combined fentanyl and THC exposure, providing a mechanistic framework linking drug exposure to impaired feto-maternal exchange and fetal growth restriction. From a translational perspective, understanding these placental mechanisms may help guide strategies to improve maternal–fetal care in populations affected by opioid use disorder and polysubstance exposure. Identifying early placental biomarkers of dysfunction, as well as pathways that could be therapeutically supported or monitored during pregnancy, may ultimately aid in risk stratification, targeted prenatal interventions, and improved surveillance of fetal growth. In the longer term, such insights may help inform clinical strategies aimed at mitigating the intergenerational health impacts of prenatal substance exposure for both pregnant individuals and their offspring.

## Acknowledgments

This work was supported by a grant from the Canadian Institutes of Health Research (CIHR) to S.B. and L.G. (grant number PJT178200). Y.C. was supported by the Brain &

Behavior Research Foundation (BBRF) Young Investigator Award (Grant number 33614) and Molly Towell Perinatal Research Fellowship. The authors acknowledge the technical support provided by the Louise Pelletier Histology Core Facility (RRID: SCR_021737), Department of Pathology and Laboratory Medicine, University of Ottawa, as well as the uOttawa Animal Behaviour and Physiology Core (RRID: SCR_022882) for assistance with indirect calorimetry experiments. The authors acknowledge the Electron Microscopy Core (RRID: SCR_025398), funded by the University of Ottawa Brain-Heart Interconnectome (BHI) through the Canadian First Research Excellence Fund (CFREF), for imaging support. The authors acknowledge the University of Ottawa Animal Care and Veterinary Service for animal housing and veterinary care.

---

**Supplementary Figure 1.**
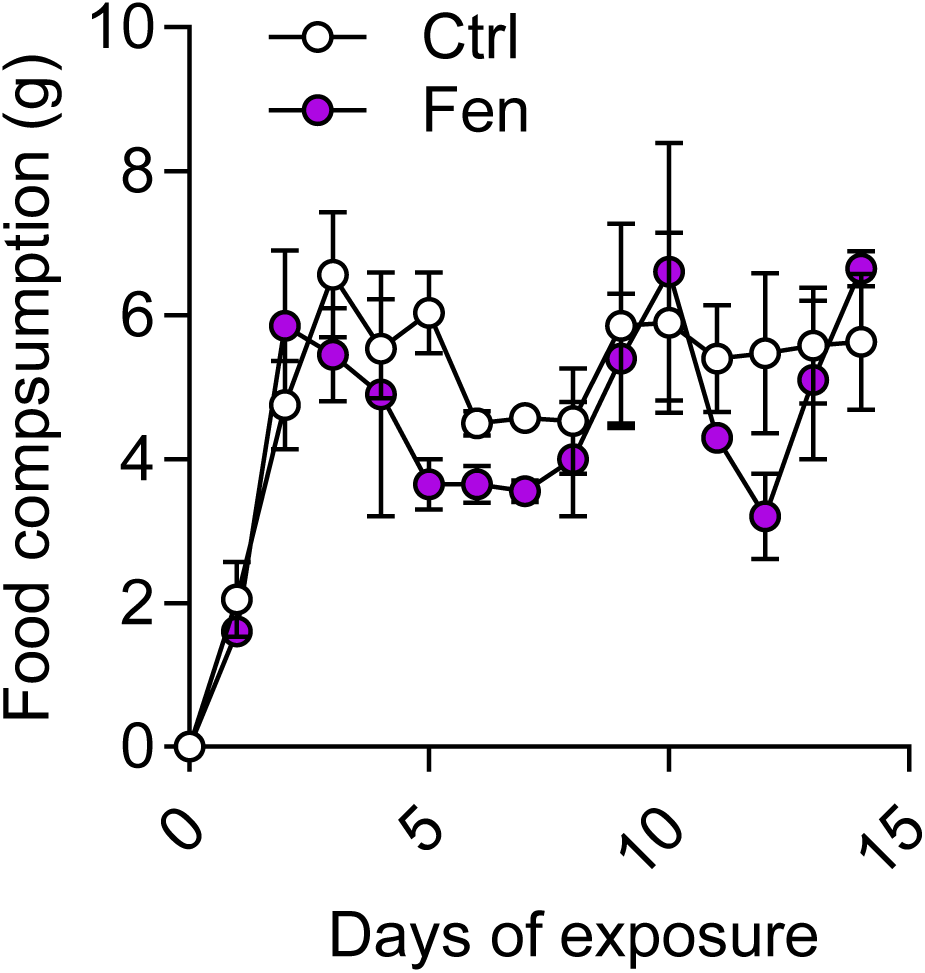
Daily food consumption during the pre-pregnancy treatment period. Food intake was recorded daily throughout the pre-pregnancy treatment period. Data are presented as mean ± SEM for each experimental group at each treatment day.

**Supplementary figure 2.**
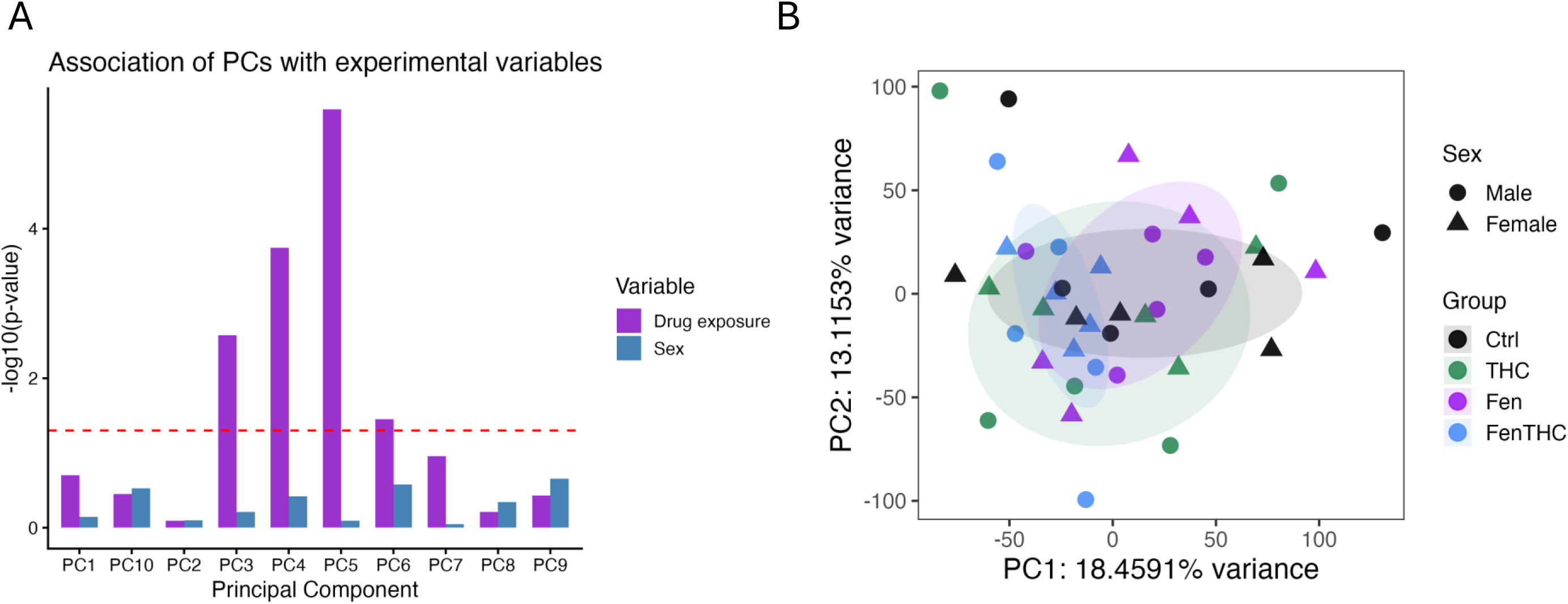
Principal component analysis of placental transcriptomic variation by exposure group and fetal sex. (A) Associations between individual principal components and experimental variables are shown as −log10(P values) for drug exposure and fetal sex. The red dashed line indicates the nominal significance threshold of P = 0.05. (B) PCA plot of placental samples using PC1 and PC2, which explain 18.46% and 13.12% of the variance, respectively. Points are colored by exposure group and shaped by fetal sex; shaded ellipses indicate group-level sample dispersion.

**Supplementary figure 3.**
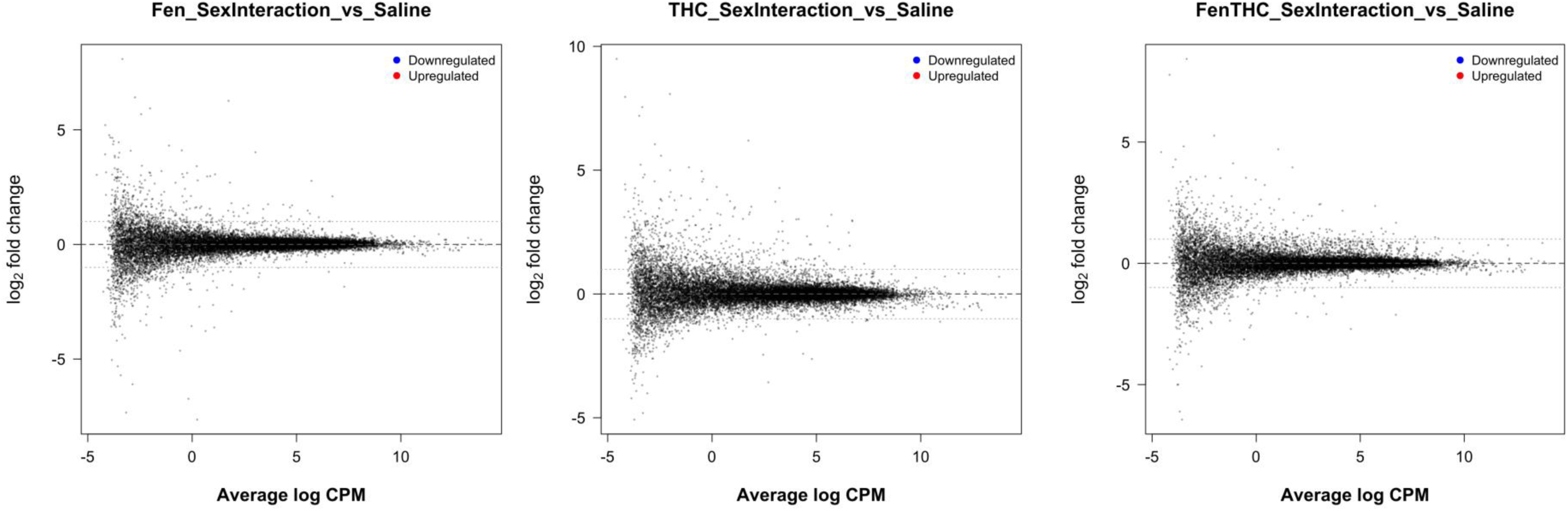
Sex-by-exposure interaction effects on placental gene expression. Mean-difference/MD plots showing sex-specific transcriptional responses to drug exposure relative to Saline controls. Each point represents one gene. The x-axis shows the average expression level as log counts per million (log CPM), and the y-axis shows the interaction log2 fold change from the edgeR model including Drugexposure × Sex. For each panel, the plotted contrast tests whether the exposure-associated change differs between female and male placentas relative to the sex difference observed in Saline controls. Positive log2 fold-change values indicate genes for which the exposure response is more positive, or less negative, in females than in males. Negative values indicate genes for which the exposure response is more positive, or less negative, in males than in females. Significantly upregulated interaction effects would be shown in red, significantly downregulated interaction effects would be shown in blue, and non-significant genes are shown in grey/black. Panels show the indicated sex-by-exposure interaction contrasts for Fen, THC, and FenTHC versus Saline.

